# Coevolution of Species’ Range Borders: Interactions Between Interspecific Competition, Gene Flow, and Matching Habitat Choice

**DOI:** 10.64898/2026.05.03.722457

**Authors:** Farshad Shirani, Judith R. Miller, Benjamin G. Freeman

## Abstract

Existing theory examining the coevolutionary dynamics of species’ range borders assumes random dispersal, which causes maladaptive gene flow from the range core to the range margins and contributes to the formation of range limits. However, dispersal is unlikely to be random for many organisms in nature, calling into question existing theoretical predictions. For example, if individuals exhibit phenotype-dependent adaptive dispersal strategies such as matching habitat choice, then the resulting adaptive gene flow toward species’ range margins could facilitate range expansions and potentially prevent the formation of range limits by interspecific competition. To test this idea, we use a comprehensive mathematical model to develop a quantitative theory of range border coevolution that incorporates phenotype-optimal dispersal—a particular form of matching habitat choice in which individuals follow the gradient in an environmental optimum phenotype to settle in the habit best suited for their phenotype. We find that instead of preventing competitively formed range limits, adaptive dispersal leads to sharper range limits and reduced character displacement in sympatry. These differences are particularly remarkable when natural selection is weak, when individuals are specialized in their resource use, or when individuals are highly sensitive to environmental conditions. We show that matching habitat choice causes backward edge-to-core movements which dynamically interact with the effects of interspecific competition to establish the range limits. Thus, the formation of range limits by interspecific competition is robust to assumptions about individual dispersal. Further, our results identify the competitive advantage of evolving matching habitat choice in steep environmental gradients, especially for slowly-growing species in rapidly fluctuating climates.

## 1. Introduction

Understanding the eco-evolutionary processes involved in the evolution of range limits between ecologically similar species is necessary to understand the establishment, structure, and stability of biological communities (Case et al., 2005; Edwards et al., 2018; Urban et al., 2008). The knowledge gained improves our ability to predict how communities will respond to climate change, and enhances management strategies for controlling invasive species (Duputié et al., 2012; Holt and Keitt, 2005; Norberg et al., 2012; Pecl et al., 2017). In principle, every biotic and abiotic factor that affects individuals’ dispersal, adaptation, reproduction, and survival, as well as their interactions with each other and with their environment, will contribute to the evolution of the species’ geographic range and borders (Angert et al., 2020; Mauro et al., 2021; Miller et al., 2020; Sexton et al., 2009; Wisz et al., 2013). A major goal of research has thus been to identify the “key” contributing factors, the presence or absence of which dramatically affects the formation and stability of the range limits. Using a comprehensive mathematical model, we present a theory explaining how interspecific competition interacts with individual dispersal and gene flow to establish borders between two species.

The role of interspecific competition in setting species’ range limits has been investigated for a long time, both empirically (Connell, 1961, 1983; Diamond, 1985; Freeman et al., 2022, 2024; Louthan et al., 2015; Pigot and Tobias, 2013; Schoener, 1983) and theoretically (Case and Taper, 2000; Case et al., 2005; Engen et al., 2021; Goldberg and Lande, 2006; Price and Kirkpatrick, 2009; Shirani and Freeman, 2025; Shirani and Miller, 2022). The work of Case and Taper (2000) stands as a major theoretical contribution. By assuming that species disperse randomly and evolve through natural selection acting on a quantitative trait, they developed a mathematical model that shows how interactions between interspecific competition and maladaptive effects of random gene flow enable the evolution of borders between two species. Although Case and Taper’s results should in general be regarded cautiously—due to a “tenuous” assumption (as they describe it) that they make on species’ trait variance—the main mechanism they describe for coevolution of species’ borders has been reaffirmed by Shirani and Miller (2022) in the absence of any assumption on trait variance.

Case and Taper’s theory relies on the assumption that dispersal is random, and hence creates maladaptive gene flow to range margins (Bonte et al., 2012; Lenormand, 2002). However, there is empirical evidence confirming the evolution of adaptive dispersal strategies in a variety of species. The individuals of such species acquire information from their environment to move towards preferred habitat (Clobert et al., 2009; Lowe and McPeek, 2014; Ponchon and Travis, 2022; Ronce, 2007; Saastamoinen et al., 2018). A rather idealized form of such biased movements is *matching habitat choice* (Box 1). The quantitative results by Shirani and Miller (2025) show that matching habitat choice causes the total gene flow to become adaptive, in both central and peripheral populations. Therefore, when species disperse adaptively, Case and Taper’s theory fails to explain whether or not interspecific competition can still result in the evolution of range borders. The uncertainty in the predictions of Case and Taper’s work is amplified by knowing that the analyses performed for a solitary species have revealed that matching habitat choice substantially enhances range expansion capacity and chance of survival when environmental gradients are exceedingly steep (Shirani and Miller, 2025). See Section S3 in the supplementary file for a summary of key predictions on range evolution of a solitary species. The main goal of the present work is to extend the theory to address these existing uncertainties and describe the eco-evolutionary mechanisms of the formation of range borders under matching habitat choice.

### Box 1: Matching habitat choice: Concepts and implications

*Matching habitat choice* is a phenotype-dependent adaptive dispersal strategy in which individuals assess their environment and move to the location that best matches their phenotype (Edelaar et al., 2008; Ravigné et al., 2004). In principle, matching habitat choice results in rapid adaptation, reduced within-population (local) genetic variation, and increased between-population genetic divergence. Importantly, matching habitat choice creates directed gene flow that compensates for, or even reverses, the maladaptive effects of random gene flow created by random movements (e.g., to explore the habitat or avoid kin competition) (Edelaar and Bolnick, 2012, 2019; Felsenstein, 1976; Garant et al., 2005; Holt, 1987; Jacob et al., 2017).

**A performance-based strategy:** Matching habitat choice spatially assorts individuals to minimize their phenotype-environment mismatch. Thus, matching habitat choice is in fact a utility- or performance-based strategy (Edelaar et al., 2008; Munar-Delgado et al., 2024; Ravigné et al., 2004). Although maximizing individuals’ performance likely increases their fitness and overall population fitness as well, it is important to note that matching habitat choice is not in principle a fitness-maximizing (ideal free) dispersal strategy. Many factors that affect population fitness, such as competition, do not directly contribute to individuals’ search for a phenotype matching environment (Shirani and Miller, 2025).

**Dependence on niche breadth:** An individual’s performance in a local environment depends on how close the conditions of the environment are to the conditions at the core of the individual’s niche (Holt, 2009). Further, it is reasonable to imagine that adaptively dispersing individuals have a fairly accurate perception of their own niche breadth. Thus, the operation of matching habitat choice depends strongly on individuals’ niche breadth, that is, the broadness of the range of resources the individuals can utilize or the environmental conditions they can tolerate. The narrower the individuals’ niche breadth, the stronger their potential to disperse to a matching habitat.

**(In)dependence on natural selection:** Even though natural selection is involved in the evolution of matching habitat choice, the operation of matching habitat choice is not driven by natural selection (Edelaar et al., 2023). In fact, it is unlikely that an individual would be able to develop a perception of the strength of natural selection in order to decide whether or not it should move to another place.

**Conflicting effects with competition:** Matching habitat choice and competition have conflicting effects on adaptation and phenotypic variation. Phenotype-dependent competition between individuals creates a frequency-dependent diversifying (disruptive) selection, which in general tends to increase local trait variance. When individuals specialize in utilizing resources based on their phenotype, the farther the phenotypes are from each other the less intense the average level of competition is within the population, hence the less loss of population fitness (growth rate). By contrast, matching habitat choice tends to spatially assort the phenotypes, bringing close phenotypes close together. This substantially reduces local trait variation and intensifies competition between phenotypes, which can then compromise the adaptive effects of matching habitat choice on population fitness.

Phenotype-dependent competition and matching habitat choice have conflicting effects on adaptation and phenotypic variation (Box 1), which makes predicting their joint contribution to range evolution a challenging problem. Including further the effects of natural selection, density-dependent effects of gene flow, and interactions with environmental gradients, makes the resulting coevolutionary range dynamics hard to predict intuitively. The mathematical framework of the model we use in the present work allows for quantifying each such effect, thereby providing a mechanistic explanation of how they interact to form range borders. The model, built on the models developed by Shirani and Miller (2022, 2025) (which are based on the seminal works of Case and Taper (2000) and Kirkpatrick and Barton (1997)), includes all the ingredients necessary to establish a quantitative theory: an environmental gradient in trait optimum, density-dependent population growth, Allee effect, directional and stabilizing selection, mutational changes in frequency of the phenotypes, phenotype-dependent competition, random dispersal, and *phenotype-optimal dispersal* —a special and possibly more realistic type of matching habitat choice (Box 2).

We develop quantitative results that explain how range borders evolve through the interactions between interspecific competition, matching habitat choice, gene flow, environmental gradients, and natural selection. We show that directed movements and the adaptive gene flow they create make the mechanism of border formation essentially different from the case where dispersal is only random. We explore the effects of key parameters such as strength of matching habitat choice, steepness of the environmental gradient, strength of natural selection, and degree of individuals’ specialization on the range borders formed between the species. Specifically, we show how these parameters affect the sharpness of the range borders and the extent of character displacement that the species exhibit where they overlap. We finally demonstrate how an adaptive dispersal strategy such as matching habitat choice confers a competitive advantage upon a species, particularly in rapidly and frequently fluctuating environments.

## 2. Methods

Our theoretical discussions are based on the results we obtain by numerically solving the equations of our mathematical model. The model is composed of a system of partial differential equations, representing the joint evolution of the population density and the mean and variance of a quantitative phenotypic trait for each of the species present in a community of competitively interacting species. The mathematical description of the model is provided entirely in Section S1 of the supplementary material. Below, we only provide a brief overview of the structure of the model. Throughout the present work, we only focus on coevolution of range borders between two species, even though the equations of the model in Section S1 are presented for the general case of a community with an arbitrary number of species. In presenting the mathematical formulations, we follow the notations used by Shirani and Miller (2022, 2025). To prevent potential misunderstandings, in Table S2 we provide a list of notational differences compared to previous models in the literature.

Individuals of each species *i, i* = 1, 2, in the model are represented by a fitness-related quantitative phenotypic trait, which is a random variable taking any value in ℝ. The model has a quantitative phenotypic framework. The populations are assumed to be fully polymorphic, meaning that all phenotype values are assumed to be present in the populations at all times. The main equations of the model (S1)–(S3) along with their nonlinear terms defined by (S4)–(S14) represent the joint evolution of species’ population density, *n*_*i*_(*x, t*), *i* = 1, 2, their trait mean *q*_*i*_(*x, t*), and their trait variance *v*_*i*_(*x, t*), where *x* denotes a point in the geographic space and *t* denotes an instance of time. For populations with normally distributed phenotypes, as we assume in our model (Assumption (iv) in Section S1.4)), the variables *n*_*i*_, *q*_*i*_, *v*_*i*_ essentially capture all that is needed to represent the populations dynamics. The parameters of the model and their range of values are given in Table S1. The choices of units for the quantities involved in the model are described in Section S1.3. The units of time, space, trait values, and population abundances are denoted by T, X, Q, and N, respectively. The default value R_*i*_ = 1 T^−1^ chosen for the maximum intrinsic growth rate of the populations in our simulations represents slowly-growing species such as birds (Niel and Lebreton, 2005), for which the evolution of matching habitat choice is likely more beneficial (Shirani and Miller, 2025).

### Box 2: Matching habitat choice: A continuum model

Following Shirani and Miller (2025), we use the term *phenotype-optimal dispersal* to refer to a special type of matching habitat choice in which individuals follow the gradient in trait optimum to eventually settle in a habitat location that minimizes their phenotype-environment mismatch. We use the continuum model of phenotype-optimal dispersal proposed by Shirani and Miller to incorporate matching habitat choice into the model of competitively interacting species we use in our study.

**Phenotypic potential energy:** To model the force perceived by individuals to disperse to matching habitats, Shirani and Miller define a *phenotypic potential energy* function, a quantitative measure of individuals’ self-assessment of (mis)performance. Denoted by *θ*_*i*_(*x, p*), the phenotypic potential energy of phenotype *p* in the *i*th species at geographic location *x* is defined by the following simplified yet meaningful function,

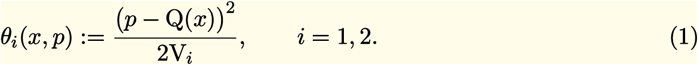

This scalar potential energy function provides a quantitative measure of how poorly the phenotype *p* is adapted to (can perform at) location *x*. Or, equivalently, *θ*_*i*_(*x, p*) determines how poorly the phenotype *p* can utilize the resources that are most favorable for phenotypes with the optimum value Q(*x*). Consistent with the concepts of matching habitat choice described in Box 1, this phenotypic potential is independent of the strength of natural selection. It depends on the degree of individuals’ phenotype-environment mismatch and the within-phenotype component of the species’ niche breadth. The latter is quantified by the variance of the individuals’ phenotype/resource utilization distributions V_*i*_ (Ackerly and Cornwell, 2007; Shirani and Miller, 2022; Violle and Jiang, 2009). See supplementary Section S1.5 for further details.

**Phenotypic dispersal force:** The phenotypic potential energy perceived by the individuals induces a *phenotypic dispersal force*,

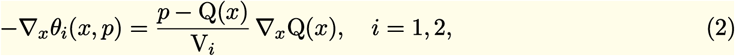

which directs the movements of phenotype *p* along the environmental gradient ∇_*x*_Q(*x*) towards a better-matching habitat location.

**Perceived environmental gradient:** A biological organism is unlikely to develop a perception of the environmental gradient in an exact mathematical sense. Further, the magnitude of the dispersal force perceived by an individual is unlikely to remain directly proportional to the steepness of the environmental gradient (as in (2)) when the gradient becomes increasingly steep. Therefore, when incorporating (2) in the model of phenotype-optimal dispersal, the environmental gradient ∇_*x*_Q is replaced with its perceived value 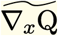. A simplified model is used for the perceived gradient such that its direction is always the same as the direction of the actual gradient but its magnitude (steepness) saturates to a maximum value when the actual gradient becomes increasingly steep. See Section S1.5 and Shirani and Miller (2025: Remark 3) for further details.

**A niche-dependent model of phenotype-optimal dispersal:** Using the dispersal force (2), the phenotype-optimal dispersal of a phenotype *p* in the *i*th species is modeled by advective (directed) movements, the velocity of which is given by A_*i*_(*x*)(−∇_*x*_*θ*_*i*_(*x, p*)), *i* = 1, 2. The parameter A_*i*_, which we assume to be constant in space (and time) throughout this work, denotes the perceived *propensity* of the individual to disperse optimally. Setting A_*i*_ = 0 in our analyses makes dispersal entirely random. Non-zero values of A_*i*_ include optimal dispersal, in addition to random dispersal which is always present in the model. The greater the value of A_*i*_ the stronger the effects of optimal dispersal.

At each location *x*, the density of phenotypes changes as a result of four different processes: random dispersal, phenotype-optimal dispersal (Box 2), intrinsic growth of the population, and mutation; see Section S1.5. The mathematical representation of these processes requires making certain assumptions, a list of which is provided in Section S1.4. The intrinsic growth rate of the phenotypes depends on the intensity of competition between them, the strength of natural selection, and Allee effect. The intensity of competition between individuals depends on how close their phenotype values are to each other, and is further adjusted by individuals’ *phenotype utilization* variance V_*i*_, *i* = 1, 2; see Section S1.5. The default value V_*i*_ = 4 Q^2^ in Table S1 results in fairly strong interspecific competition, noting that the per capita effects of competition between phenotypes follow a Gaussian form with variance 2V_*i*_ when V_*i*_ takes the same value for both species; see the competition kernel given by equation (S20). Natural selection penalizes the phenotype values that are different from the environmental optimum phenotype Q(*x*). We assume Q changes linearly in space, representing an environmental gradient ∇_*x*_Q with constant steepness.

The model includes both random and phenotype-optimal dispersal. The random component can represent the exploratory movements that individuals may initially perform to assess the environment and move to a matching location. This component can also represent any other uninformed movements to escape kin competition or avoid inbreeding. The phenotype-optimal component is modeled as described in Box 2. The strength of this dispersal component depends on individuals’ perceived *propensity* to disperse adaptively, denoted by parameter A_*i*_ for the *i*th species. Setting A_*i*_ = 0 makes dispersal entirely random. Non-zero values of A_*i*_ then include optimal dispersal, in addition to the always present random dispersal.

## 3. Results

To obtain our numerical results, we consider a one-dimensional habitat with reflecting boundaries, wherein the two populations are initially distributed allopatrically. The details of population initialization are described in Section S2.2. We assume the trait optimum Q increases linearly along the habitat. Using this simulation layout, we numerically solve equations (S1)–(S14) with parameter values specified independently for each simulation.

### 3.1 Coevolution of species’ range borders

Figure 1 shows evolutionary population dynamics of the species in a steep environmental gradient, both in the absence (left panel) and in the presence (right panel) of phenotype-optimal dispersal. When dispersal is only random, the coevolution of the borders follows the existing theory (Case and Taper, 2000; Shirani and Miller, 2022). The initial populations gradually adapt (*q*_*i*_ → Q) to new locations and expand their range. The curves of trait mean in Figure 1c show significant maladaptation near range margins, where the curves are flattened and fail to follow the slope of the gradient. This is due to the maladaptive effects of asymmetric core-to-edge gene flow. As a result, trait variance also declines at range margins (Figure 1e); see the discussions by Shirani and Miller (2022) for further details. At the time the two populations meet at the center of the habitat and become sympatric over a short region, they are both fairly well-adapted to the environment. Therefore, a strong competition is initiated between them, which induces character displacement (hence departure of *q*_1_ and *q*_2_ from Q) in their overlapping sub-populations. The resulting maladaptation along with the direct fitness loss caused by competition substantially reduces the density of the populations at their interface. As a result, the asymmetry in core-to-edge gene flow within each species is intensified, moving the species’ trait mean over the region of sympatry further away from the optimum (increasing character displacement). As range edges advance further and the spatial overlap between the species grows, this reinforcing feedback between competitively-induced character displacement and maladaptive gene flow continues to create stronger levels of maladaptation at range edges. It eventually prevents local adaptation at edges and halts range expansions, resulting in the formation of range borders (Figure 1a.)

**Figure 1:**
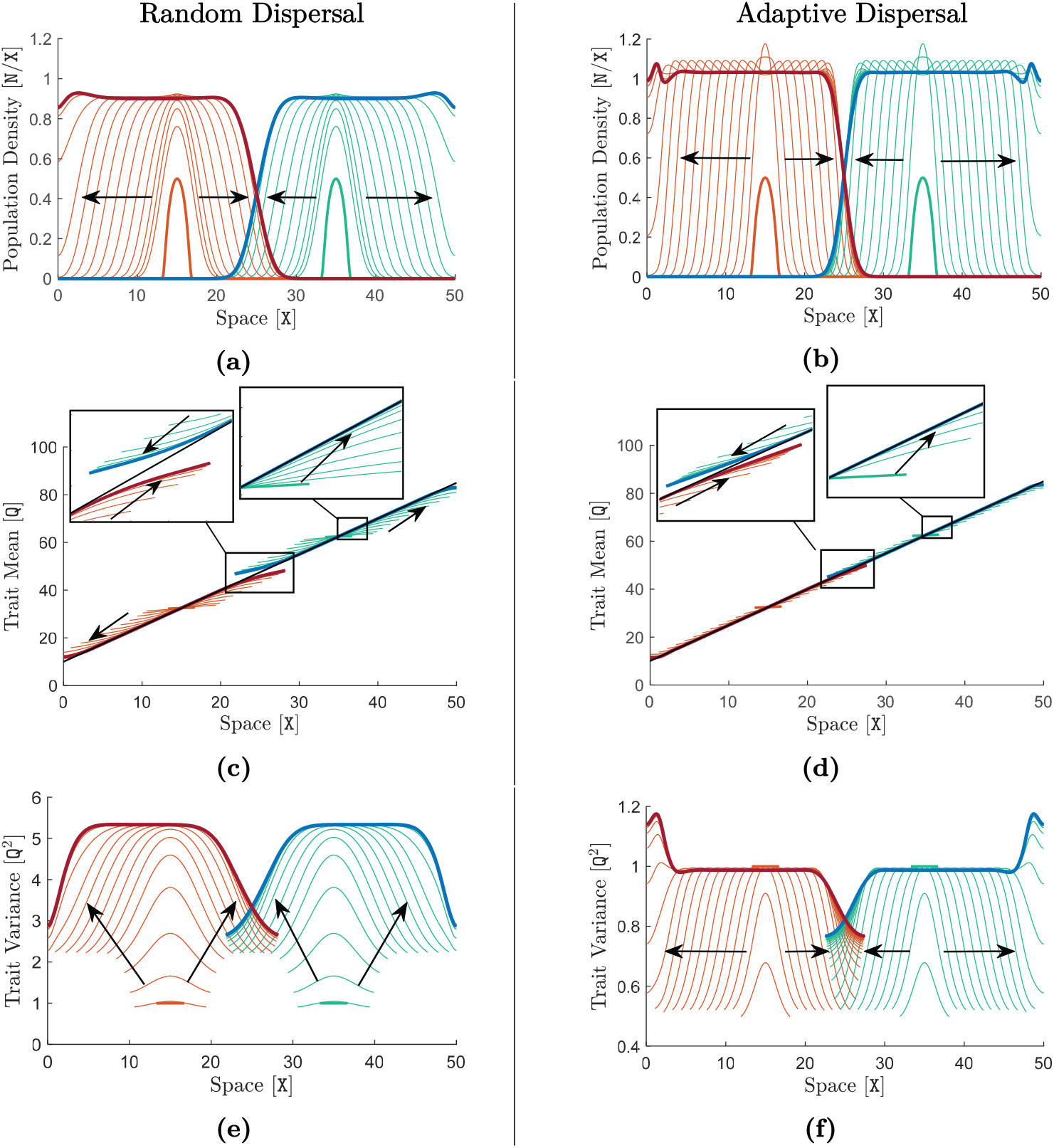
Coevolution of species’ range borders and formation of a region of sympatric coexistence. The two species simulated in the left panel (graphs (a), (c), and (e)) disperse only randomly, that is, A_1_ = A_2_ = 0 X^2^*/*T. In the right panel, the two species perform strong phenotype-optimal dispersal with A_1_ = A_2_ = 10 X^2^*/*T. The rest of the parameters of the two species in both panels are set equal to the default values given in supplementary Table S1. The trait optimum Q is linear and increasing, shown by the black line in (c) and (d), with a steep gradient of ∇_*x*_Q = 1.5 Q*/*X. Curves of the species’ population density *n*_*i*_, as their range evolves in time, are shown in (a) and (b). Curves of the species’ trait mean *q*_*i*_ are shown in (c) and (d). Curves of the species’ trait variance *v*_*i*_ are shown in (e) and (f). Note the difference in the scales of the y-axis in (e) and (f). In all graphs, curves are shown at every 4 T for a simulation time horizon of *T* = 300 T, at which an equilibrium state is approximately formed. For the 1st (left) species, curves are shown in orange, thick orange curves indicate the initial curves at *t* = 0 T, and the final curves at the end of the simulation are highlighted in red. For the 2nd (right) species, curves are shown in green, thick green curves indicate the initial curves, and the final curves are highlighted in blue. Arrows show the direction of evolution in time. Although the solutions of the model are computed for the entire habitat, the values of trait mean and trait variance outside the range of the species are not biologically meaningful. Therefore, the portions of the curves that are outside the species’ effective ranges (i.e., the regions where *n*_*i*_ *<* 0.02, *i* = 1, 2), are not shown in (c)–(f). Graphs (a) and (b) show that the species’ ranges converge to an equilibrium state, at which a limited region of sympatry is formed at the interface of the two species in the middle of the habitat. The insets in (c) and (d) highlight the formation of character displacement over the regions of sympatry, as well as the speed of species’ adaptation (convergence to the trait optimum) to the environment.

The results shown in Figure 1 confirm that range borders are also evolved in the presence of adaptive dispersal. However, compared with the random dispersal case, we observe several distinctive differences. Specifically, the region of sympatry at equilibrium is shorter and species’ borders are sharper (Figure 1b). The species’ population density presents a small overshoot near range edges, which disappears at equilibrium. Adaptation occurs much more rapidly, and the extent of character displacement at equilibrium is substantially smaller (insets in Figure 1d). Finally, the local trait variance is dramatically lower (Figure 1f). To understand the causes of these differences and the underlying mechanism of the formation of range borders, we compute the contribution of each component of dispersal to the rates of changes in species’ trait mean, ∂_*t*_*q*_*i*_, and species’ population density, ∂_*t*_*n*_*i*_, *i* = 1, 2. We also compute the local contributions to ∂_*t*_*q*_*i*_ and ∂_*t*_*n*_*i*_ due to the effects of selection and competition. The details of these computations are provided in Section S2.3. The results are shown inf Figures 2 and 3 for the 1st species and in Figure S1 for the 2nd species.

**Figure 2:**
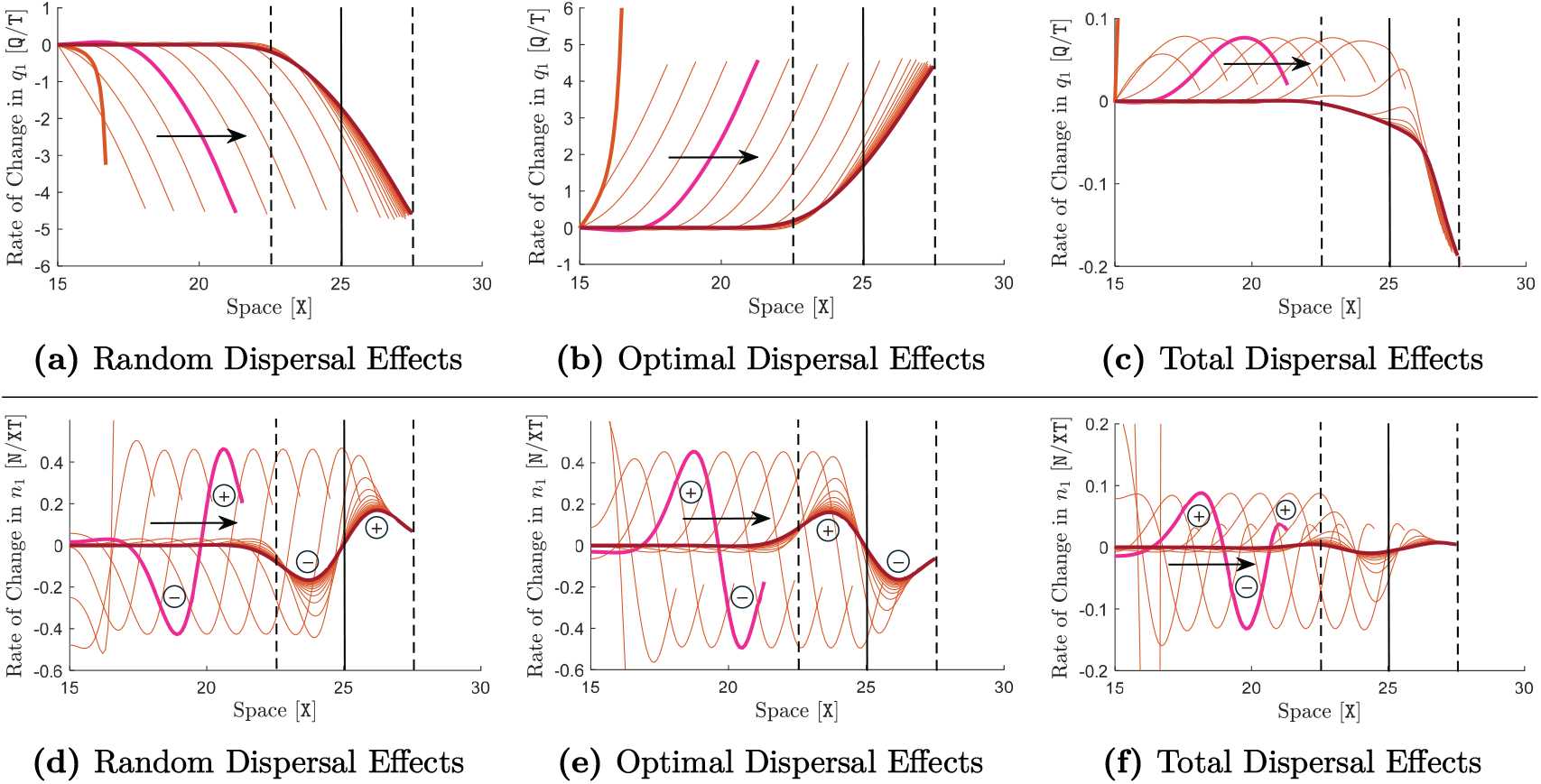
Effects of random and optimal dispersal on rates of change in trait mean and population density. The graphs shown here are associated with the same simulation performed for the results shown in the right panel of Figure 1, that is, with A_1_ = A_2_ = 10 X^2^*/*T and ∇_*x*_Q = 1.5 Q*/*X. Similar graphs but associated with the left panel of Figure 1 (A_1_ = A_2_ = 0 X^2^*/*T) are shown in supplementary Figure S2. The contributions of random dispersal (random gene flow), phenotype-optimal dispersal (directed gene flow), and total dispersal (net gene flow) to the rate of change of trait mean (adaptation or maladaptation rate) within the 1st species, ∂_*t*_*q*_1_, are shown in graphs (a)–(c) in the upper panel. The contributions of these dispersal components to the rate of change of population density of the 1st species, ∂_*t*_*n*_1_, are shown in the lower panel. Note the difference in the scales of the y-axis in all graphs. The details of the computations associated with these contributions are described in supplementary Section S2.3. The curves in all graphs are shown only for the right half of the species’ range, where it meets and coevolves with the 2nd species. The curves for the 2nd species are shown in supplementary Figure S1. In all graphs, curves are shown at every 4 T, and the portions of the curves that lie outside the species’ range are not shown. The final equilibrium curves obtained (approximately) at the end of the simulation (*T* = 300 T) are highlighted in red. The sample curves highlighted in pink are associated with the species’ range expansion regime before meeting the 2nd species. The solid black lines indicate the center of the habitat where the interface between the two species (Figure 1b) is formed. The dashed lines indicate the boundaries of the region of sympatry formed at the interface.

**Figure 3:**
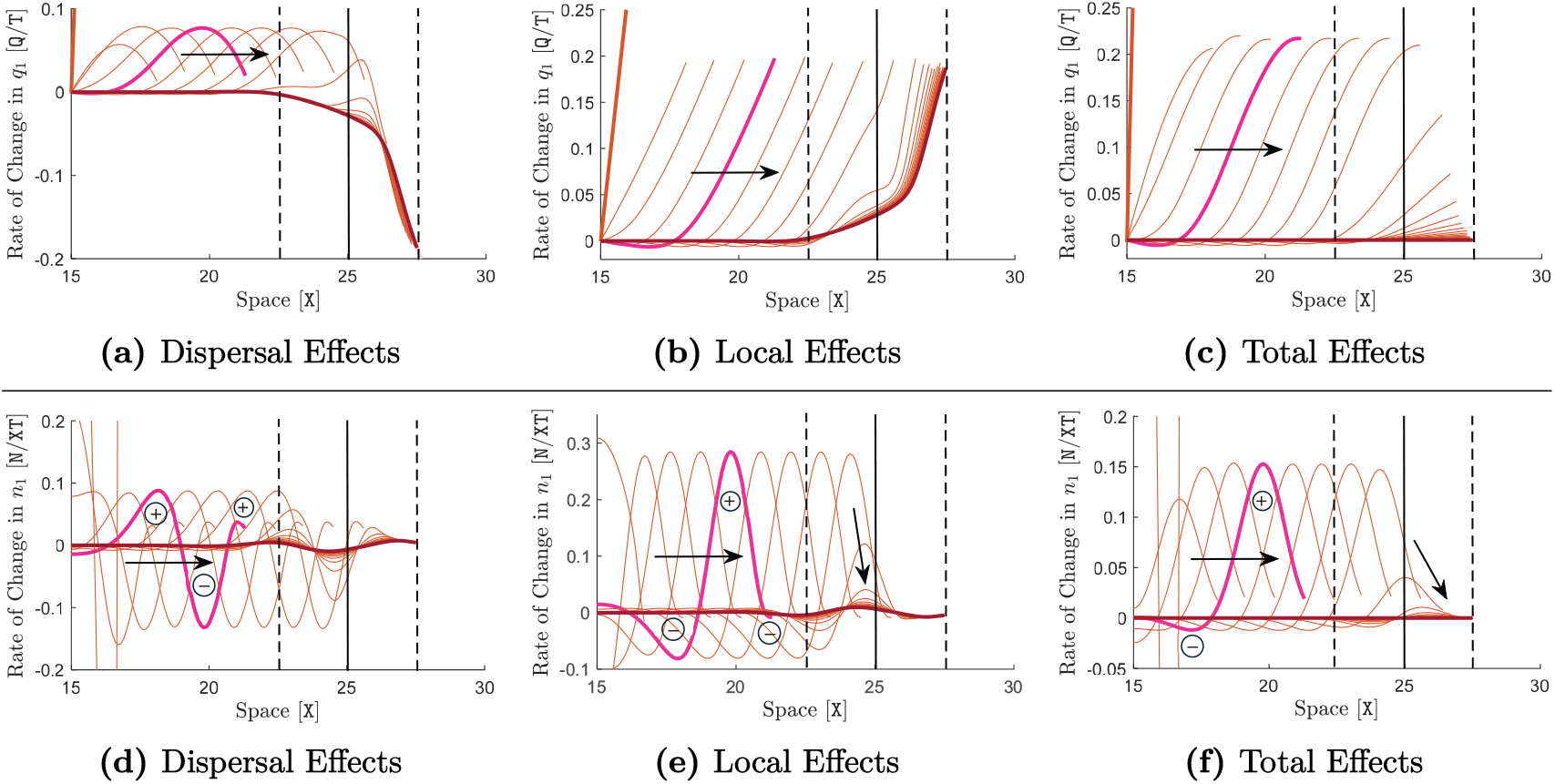
Effects of adaptive dispersal, selection, and competition on rates of change in trait mean and population density. The graphs shown here complete the set of graphs shown in Figure 2 to demonstrate the effects of different components contributing to ∂_*t*_*q*_1_ (top panel) and ∂_*t*_*n*_1_ (bottom panel). As in Figure 2, the graphs are associated with the same simulation shown in the right panel of Figure 1, that is, with adaptive dispersal. Similar graphs but associated with the left panel of Figure 1 (random-only dispersal) are shown in supplementary Figure S3. Graphs (a) and (d) are the same graphs shown in Figures 2c and 2f, which are repeated here for simplicity of comparison. They show the contribution of total adaptive dispersal (random plus phenotype-optimal) to ∂_*t*_*q*_1_ and ∂_*t*_*n*_1_, respectively. The local contributions of selection and competition to ∂_*t*_*q*_1_ and ∂_*t*_*n*_1_ are shown in (b) and (e), respectively. The details of the computations associated with these contributions are described in supplementary Section S2.3. Graphs (c) and (d) show the total effects. That is, (c) shows exactly the curves of ∂_*t*_*q*_1_, obtained as the sum of the curves in (a) and (b). Similarly, (f) shows exactly the curves of ∂_*t*_*n*_1_, obtained as the sum of the curves in (d) and (e). The same description as in Figure 2 holds for the curve colors and the solid and dashed lines. Note the difference in the scales of the y-axis in all graphs.

The contributions of the dispersal components to ∂_*t*_*q*_1_ confirm strong adaptive effects of the directed gene flow created by phenotype-optimal dispersal (Figures 2a–2c). To see these effects, first note that positive values of ∂_*t*_*q*_1_ at a location *x* imply that the species’ trait mean increases at *x*. Contrariwise, negative values of ∂_*t*_*q*_1_ imply a decrease in the trait mean. Also, note that the curves in Figure 2 are shown for the right half of the 1st species’s range, in which the initial profile of the species’ trait mean lies below the trait optimum; see Figure 1d. Therefore, positive values of ∂_*t*_*q*_1_ imply adaptive effects (increasing *q*_1_ towards Q) and negative values imply maladaptive effects. Based on these observations, Figure 2a confirms that the random gene flow created by random dispersal is always maladaptive, particularly at range margins (wavefronts)—a result consistent with the known disruptive effects of random gene flow (Kirkpatrick and Barton, 1997; Lenormand, 2002; Shirani and Freeman, 2025). By contrast, Figure 2b shows that the directed gene flow created by matching habitat choice is always adaptive, even at range margins. Further, it almost completely compensates for the maladaptive effects of random gene flow. As a result, the total gene flow caused by overall dispersal becomes adaptive, except over a short region near the range edge at equilibrium (see the red curve in Figure 2c). The creation of this adaptive gene flow explains the rapid adaptation shown in Figure 1d. It also implies that disruptive gene flow does not play a major role in forming borders when species disperse adaptively. We note that the maladaptive effects observed in Figure 2c near the range border at equilibrium are much weaker than the swamping effects observed in the absence of matching habitat choice; see Figure S2c. We note further that natural selection also acts locally to adapt the populations (Figure 3b). However, the adaptation caused by selection in the presence of phenotype-optimal dispersal is much slower than the adaptation caused when dispersal is only random (Figure S3b). This is because the phenotype-environment matching caused by phenotype-optimal dispersal substantially removes the need for selection to act.

When dispersal is only random, movements near the range margins are strongly asymmetric, predominantly from the populous center of the species to their sparsely populated periphery—hence creating the asymmetric core-to-edge gene flow we discussed before. These asymmetric movements are schematically illustrated in Figure S4a. They can be seen more accurately through the spatial profile of the contribution of dispersal to ∂_*t*_*n*_*i*_, shown for the 1st species in Figure S2f. This profile is also approximately preserved in the contribution of the random component of dispersal in the presence of optimal dispersal; see Figures 2d and S1d. At the wavefront of the species’ population density, where the density declines from its maximum (at core) to almost zero (at edge), the contribution of random dispersal to ∂_*t*_*n*_1_ shows a transition from negative values (near the core) to positive values (near the edge). This (−)(+) transition pattern can be seen in Figure S2f and is present both during the range expansion regime and at equilibrium. The transition point from (−) to (+) occurs at the inflection point on the wavefront of the curve of population density in Figure 1a. The negative values of ∂_*t*_*n*_1_ in the core side of the wavefront implies that the “overall” random movement over that region is outward, that is, towards the edge. As a result, the contribution of random dispersal to ∂_*t*_*n*_1_ becomes positive (increasing the density) near the edge. This confirms that interspecific competition and the strong migration load caused by random gene flow are then the factors which act locally (Figure S3e) to canceling out the density-increasing effects of random movements near the edge, permitting the establishment of borders at equilibrium. This cancellation can be seen through Figures S3d–S3f.

Figures 2d–2f demonstrate a key process contributing to the coevolution of range borders in the presence of matching habitat choice: backward edge-to-core movements that completely compensate for, and even partially reverse the effects of core-to-edge movements caused by random dispersal; see also the illustration in Figure S4. The overall edge-to-core movement is implied by the (+)(−) pattern in the contribution of adaptive dispersal to ∂_*t*_*n*_1_ at wavefronts (Figure 2e), which is opposite to the (−)(+) pattern of contribution of random dispersal. When the individuals at the core randomly move to or beyond the range edge, for example to explore their surroundings during range expansion, the majority of them find the new habitat less suitable for their phenotype. Matching habitat choice then pushes them to move back to the core. These backward movements at wavefronts underlie several of the distinctive differences we observed in Figure 1 in comparison with the case of random-only dispersal. During the range expansion regime, the backward movements cause the contribution of overall dispersal to ∂_*t*_*n*_1_ to follow a (+)(−)(+) pattern at wavefronts (Figure 2f). Similar to the case of random-only dispersal (Figures S3d and S3e), the local contribution of competition and selection to ∂_*t*_*n*_1_ then follows an opposite pattern (−)(+)(−); see Figures 3d and 3e. This results in the complete profile of ∂_*t*_*n*_1_ to become slightly negative near the core before transitioning to positive at the wavefronts (Figure 3f), explaining the overshoots observed in range expansion waves in Figure 1b. It is worth noting that, the patterns we see in the contributing components to ∂_*t*_*n*_1_ remain qualitatively unchanged when gene flow is predominantly directed (optimal). We made this observation by simulating an artificially strong level of phenotype-optimal dispersal with A_*i*_ = 50 X^2^*/*T, that is not shown here.

When the two species meet, interspecific competition and backward edge-to-core movements induced by matching habitat choice interact to establish the range limits (Figure S4b). Interspecific competition tends to induce character displacement over the region of sympatry gradually formed between the species. However, character displacement creates a phenotype-environment mismatch in the sympatric subpopulations. Matching habitat choice then pushes the individuals at range margins to move back to better-matching locations at the core, hence reducing the extent of character displacement. These interactions between interspecific competition and backward movements eventually reach a balanced steady state, with significantly reduced character displacement compared with the case of random-only dispersal. The evolving curves in Figures 1b, 1d, 2, and 3 confirm the presence and convergence to this equilibrium state. At equilibrium, the overall edge-to-core movement resulting from adaptive dispersal almost completely cancels out the core-to-edge random movements, leading to almost no density change by total dispersal (Figure 2f)—a dramatic difference compared with the random-only dispersal case (Figure S2f). This also explains the enhanced sharpness of the borders in the presence of adaptive dispersal (Figure 1b). Since overall dispersal does not significantly contribute to density change at equilibrium, the density decline from core to edge near the borders is predominantly due to intensified interspecific competition that acts locally in space, as well as the slight maladaptive gene flow that is created only close to range edges at equilibrium (Figure 2c).

### 3.2 Effects of dispersal, environment, and specialization on character displacement and sharpness of the borders

We explore further the differences between the coevolution of borders under matching habitat choice and under random dispersal, by computing the extent of character displacement and the length of the region of sympatry at equilibrium. We perform computations for random-only and moderate and strong optimal dispersal propensity, as well as broad ranges of values for the steepness of the environmental gradient, the strength of stabilization selection, and the individuals’ specialization level. The results are shown in Figure 4.

**Figure 4:**
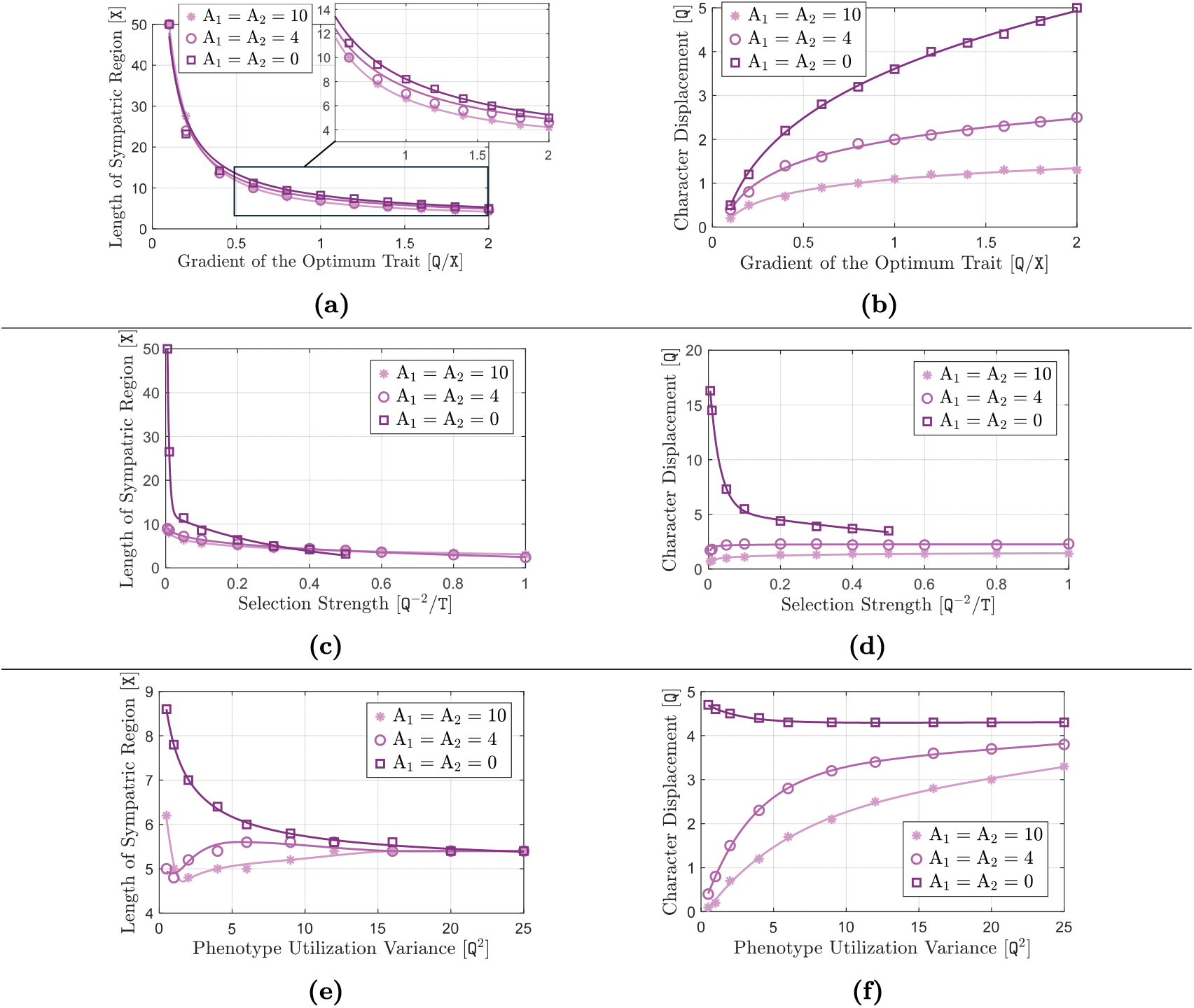
Effects of matching habitat choice, gradient steepness, selection strength, and individuals’ specialization on character displacement and region of sympatry at equilibrium. The lengths of the region of sympatry when species’ ranges converge to an equilibrium state (e.g., as in Figures 1c and 1d) are shown in (a), (c), and (e). Correspondingly, the extents of character displacement at equilibrium are shown in (b), (d), and (f). In the upper, middle, and lower panels, the results are shown, respectively, for a broad range of values of the steepness of the environmental gradient, the strength of the stabilizing selection (S), and the variance of the phenotype (resource) utilization distribution (assuming V_1_ = V_2_). In all graphs, the results are shown for three different values of the optimal dispersal propensity, with A_1_ = A_2_ = 0 X^2^*/*T resulting in only random dispersal. The environmental gradient for the results shown in the middle and lower panels is set to be steep, ∇_*x*_Q = 1.5 Q*/*X. The rest of the parameters take their default values given in Table S1. The data points are obtained as follows. At different values of the variable parameter, a simulation similar to those associated with Figure 1 is performed for a period of time long enough for species’ range evolution to converge to a steady-state (equilibrium). In each simulation, the region of sympatry is identified as the region over which both species coexist with a density greater than 0.02 N*/*X. The lengths of the identified regions are marked by the data points in (a), (c), and (e). Denoting the equilibrium trait mean of the *i*th species by 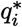, the extent of character displacement is computed as the maximum value of 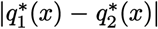 when *x* takes all values in the region of sympatry. In our simulations, the maximum character displacement always occurs at the species’ boundary. The computed character displacements are marked by the data points in (b), (d), and (f). The curves are obtained by interpolating the data points. We note that, in the absence of optimal dispersal (A_1_ = A_2_ = 0), the species’ range expansion becomes too slow when S takes values greater than 0.5 Q^−2^*/*T. In this case, the species eventually become extinct when S ≳ 0.88 Q^−2^*/*T, the critical value that can be computed using the formula given by Shirani and Miller (2022: Remark 4).

Figure 4a shows that the region of sympatry expands when the environmental gradient becomes shallower. When the gradient is sufficiently shallow (∇_*x*_Q → 0), the species become sympatric over the entire habit and no borders are formed between them. This is because in shallow gradients the maladaptive core-to-edge effects of random dispersal do not become strong enough to destabilize the competition-selection balance at range margins and set range limits. The state of full sympatry evolves as ∇_*x*_Q → 0 regardless of the strength of optimal dispersal propensity. In fact, in shallow gradients the total dispersal is almost random even if A_*i*_ takes large values—letting ∇_*x*_Q → 0 in (2) immediately implies that the optimal dispersal force vanishes to zero. It is also fairly intuitive to imagine that in shallow gradients individuals do not attain a sufficiently strong phenotype-environment match by moving along the gradient. In steep gradients, however, the backward movements induced by strong adaptive dispersal reduce the span of the region of sympatry, as described before.

Figure 4b shows that character displacement increases as the environmental gradient becomes steeper. In steeper gradients the maladaptive effects of random gene flow to range margins are stronger, and hence it enhances further the character displacement induced by interspecific competition. However, the rate of increase in character displacement as the gradient becomes steeper is much lower in the presence of adaptive dispersal. As we described before, in steep gradients matching habitat choice effectively pushes back the peripheral populations whose phenotypes are significantly displaced from the optimum.

When stabilizing natural selection is weak (S → 0), Figures 4c and 4d reveal a remarkable difference between the borders formed under matching habitat choice and those formed under random dispersal. When dispersal is only random and selection is sufficiently weak, no borders are formed between the species even though they express exceedingly strong character displacement. This is because the maladaptive effects of random gene flow are not “felt” by the species due to the weakness of the selection. Strong character displacements are expressed because only at such levels of displacement from trait optimum do the stabilizing effects of weak selection become sufficiently strong to balance the diversifying effects of competition. It is worth noting that the sharp increase in character displacement and span of the region of sympatry in Figures 4c and 4d, when A_1_ = A_2_ = 0, arises when selection strength falls approximately below 0.05 Q^−2^*/*T. Although selection at this level is considered weak, it is still stronger than almost 25 percent of the estimates provided by Shirani and Miller (2022: Fig. 1) based on the data available for 62 different species (Kingsolver et al., 2001, 2008). By contrast, range borders are formed under matching habitat choice even when selection is very weak. The region of sympatry expands only slightly with weak selection. The extent of character displacement is almost insensitive to selection strength and remains significantly reduced. The reason behind these distinct differences is that, as we discussed in Section 3.1, maladaptive gene flow does not play a major role in coevolution of borders under matching habitat choice. Thus, the evolution of range limits in this case is rather insensitive to the strength of natural selection.

Another remarkable difference emerges for species with highly specialized individuals (V_*i*_ → 0, *i* = 1, 2). Figure 4e shows that, with random-only dispersal, the length of the region of sympatry increases significantly when individuals become exceedingly specialized. This is because specialized populations effectively release themselves from competition. Hence, a broader region of sympatry is formed before the level of character displacement is sufficiently enhanced by maladaptive gene flow to halt range expansions. By contrast, with adaptive dispersal the length of the region of sympatry is fairly insensitive to changes in the degree of specialization. This is because matching habitat choice strongly controls the level of local trait variation (Figure 1f), keeping the specialized individuals sufficiently close together to still engage in strong competition. Figure 4f shows that the extents of character displacement are also remarkably different in different dispersal cases. With random-only dispersal, character displacement is fairly insensitive to the specialization level. In stark contrast, excessive specialization substantially reduces character displacement by dramatically strengthening the optimal dispersal force (see equation (2)) and hence the backward movements it induces near range borders.

### 3.3 Competitive advantages of matching habitat choice in steady and rapidly fluctuating environments

We conclude our results by showing that, in steep environments, evolution of matching habitat choice in a species can give the species significant competitive advantage over a randomly dispersing similar species. The advantage is particularly notable for slowly-growing (small R_*i*_) species in rapidly fluctuating environments, for example, species of montane birds living on temperate mountains with drastic seasonal variations in climate. We perform simulations of two species with identical parameters, but one dispersing through matching habitat choice with a moderate dispersal propensity A_1_ = 2 X^2^*/*T, and the other dispersing only randomly. We consider both a steady environment (fixed Q) and a periodically fluctuating environment. The results are shown in Figure 5.

**Figure 5:**
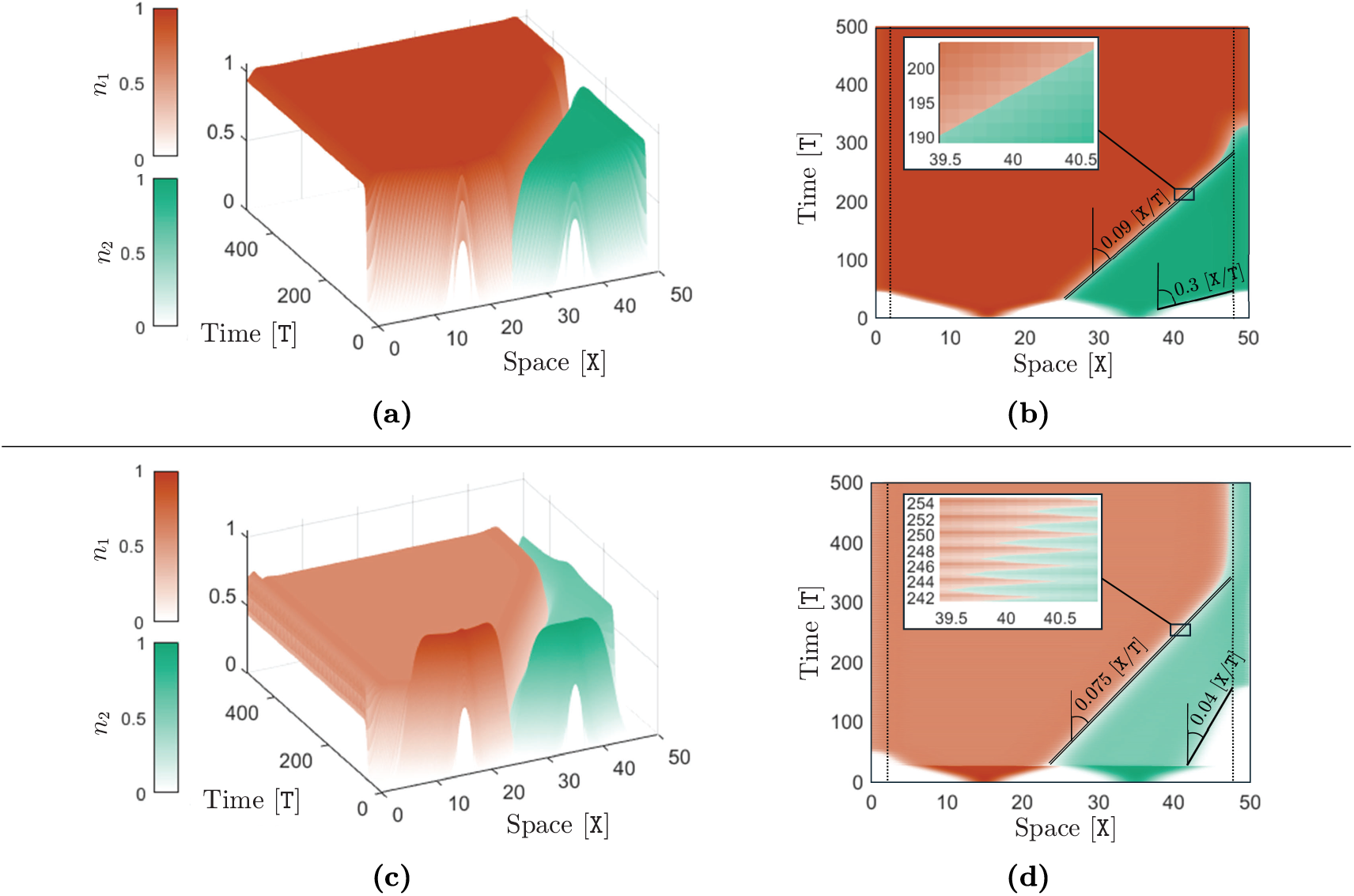
Competitive advantage of matching habitat choice in steady and rapidly fluctuating environments. The 1st species expresses a moderate level of phenotype-optimal dispersal with A_1_ = 2 X^2^*/*T whereas the 2nd species disperses only randomly, A_2_ = 0 X^2^*/*T. The environmental gradient is steep, ∇_*x*_Q = 1.5 Q*/*X, and the rest of model parameters take their default values given in Table S1. The graphs (a) and (b) in the upper panel show the results of a simulation with a steady (temporally constant) environment, whereas the graphs (c) and (d) in the lower panel are associated with a periodically fluctuating environment. The evolution of the species’ population density *n*_*i*_, *i* = 1, 2, is shown in each graph. Unlike Figure 1, the time axis is shown explicitly to allow for clear visualization of the moving borders. The graphs (b) and (d) show the top view of the same graphs shown in (a) and (c), respectively. The slopes specified on graphs (b) and (d) give the speed of the moving borders formed between the species (double line) as well as the other moving edge of the 2nd (competitively weaker) species. The dotted lines in (b) and (d) indicate the boundaries of the region Ω_*δ*_ as described in the discussion following equation (S14) in the supplementary material. It is assumed that outside Ω_*δ*_ (near the habitat boundaries) the individuals change their optimal dispersal behavior to avoid crossing the habitat boundaries. In the lower panel, the temporal fluctuations in the environment start at *t* = 30T—when borders have effectively been established between the species—and follow a square waveform with period 2 T. Specifically, at the beginning of each period the (linear) spatial profile of the trait optimum Q is abruptly shifted up by 4 Q and then remains fixed (at the shifted profile) for the first half of the period (for 1 T). It is then shifted down by the same amplitude 4 Q back to the initial profile and remains fixed (at the initial profile) for the second half of the period (for 1 T). This fluctuation pattern is repeated periodically until the end of the simulation.

The evolution of the species’ population density, shown for a steady environment in Figures 5a and 5b, confirms the competitive advantage of the species performing matching habitat choice. The borders formed between the species at the middle of the habitat are constantly pushed towards the randomly dispersing species, leading to exclusion of this species from the habitat. This competitive advantage of adaptive dispersal is particularly strong when the environment fluctuates rapidly and frequently. To see this, we simulate periodic abrupt fluctuations in the trait optimum, with period 2 T and amplitude 4 Q. At the beginning of each period the trait optimum Q is abruptly shifted up by 4 Q and then remains fixed for the first half of the period. It is then shifted down by 4 Q back to its initial value and remains fixed for the second half of the period. The results are shown in Figures 5c and 5d. We first note that frequent environmental fluctuations dramatically reduce range expansion speed of a randomly dispersing species. This is confirmed by comparing the speed 0.3 X*/*T of the freely moving edge of the randomly dispersing species in the steady environment (Figure 5b) with its speed of 0.04 X*/*T in the fluctuating environment (Figure 5d). By contrast, comparing the speeds 0.09 X*/*T and 0.075 X*/*T of the moving borders in the steady and fluctuating environments, respectively, indicates only a slight reduction due to the fluctuations. This further implies that the range expansion speed of the adaptively dispersing species is much less sensitive to environmental fluctuations. If the dispersal propensity of this species is made stronger, fluctuations are made more frequent, and habitat is made wider, then the randomly dispersing species may become extinct even before reaching the habitat boundary.

## 4. Discussion

We find that predictions of models demonstrating the formation of range limits driven by inter-specific competition are robust to assumptions about dispersal. Previous theory on competitively formed range limits (Case and Taper, 2000; Shirani and Miller, 2022) assumed that species disperse randomly, with the maladaptive effects of core-to-edge gene flow being a key contributor to the formation of range limits. This theory failed to address a situation where species disperse adaptively, which leads to peripheral populations receiving adaptive gene flow from core populations (Shirani and Miller, 2025). It thus seemed plausible that incorporating adaptive dispersal into models could lead to different outcomes on the possibility of range limits to be set by interspecific competition. However, this is not what we found. Instead, including matching habitat choice in models led to sharper range borders and reduced character displacement compared to models assuming random dispersal.

We showed that, when species disperse adaptively, the key factor that contributes along with interspecific competition to formation of range limits—despite gene flow being adaptive—is the backward edge-to-core movement caused by adaptive dispersal at species’ range expansion wave-fronts. The individuals which move randomly to expanding range margins, for example to explore new habitats or avoid kin competition, often face environments less suited to their phenotype. Following their preference for a matching habitat, they then move back towards the range core. When two competing species meet, these backward edge-to-core movements act to reduce the extent of character displacement induced by interspecific competition over the region of sympatry between the species. At equilibrium, the backward movements effectively cancel out the effects of random core-to-edge movements, resulting in almost no density change due to overall movements of individuals at and behind range borders. Interspecific competition, intensified further by reduced character displacement, then acts to sharply reduce species’ population density and set range limits.

The effects of matching habitat choice on increasing the sharpness of the borders and reducing character displacement become remarkably pronounced when stabilizing natural selection is weak or when individuals have a narrow niche breadth; see Figures 4c–4f and Boxes 1 and 2. When dispersal is random, no borders may form in a linearly changing environment if the environment is weakly selective. That is, the species may become sympatric all over the habitat. Under matching habitat choice, by contrast, range borders are formed and remain relatively sharp, and species express reduced character displacement regardless of the strength of natural selection. When individuals’ niche breadth is narrow, that is, when individuals are highly specialized in utilizing resources or are very sensitive to environmental conditions, matching habitat choice dramatically reduces character displacement. However, we should note that the effects of matching habitat choice are prominent only if the environmental gradient is steep; see Figures 4a and 4b. In fact, considering its high cost (Bonte et al., 2012; Bowler and Benton, 2005; Clobert et al., 2009; Travis et al., 2012), matching habitat choice is unlikely to evolve as a dispersal strategy in shallow gradients (Shirani and Miller, 2025).

Matching habitat choice is hard to detect in nature and empirical evidence for its presence in nature is still limited (Camacho et al., 2020; Edelaar and Bolnick, 2012, 2019; Edelaar et al., 2023; Shirani and Miller, 2025). Our results suggest that the distinctive characteristics of range borders in the absence of sufficiently strong natural selection—that is, strikingly lower degrees of character displacement and sharper range borders—may be indications of matching habitat choice. The sharpness of the borders should be measured in our general choice of the unit of space (see Section S1.3) and compared with species of similar characteristic parameters in equally steep gradients. This adds to the set of other hallmarks of matching habitat choice, such as adaptive gene flow to range margins and substantially reduced trait variation at central populations (Shirani and Miller, 2025). Our results further confirm the particular advantages of matching habitat choice for slowly-growing (small values of R_*i*_) species in rapidly fluctuating environments, through enhancing the species’ competitive strength, invasion capacity, and persistence; see Figure 5.

In summary, we find that interspecific competition forms range limits across a range of assumptions about individual dispersal. That a variety of theoretical approaches generate the same general outcome (Case and Taper, 2000; Case et al., 2005; Goldberg and Lande, 2006; Price and Kirkpatrick, 2009; Roughgarden, 1979; Shirani and Miller, 2022; Tilman, 1982) is consistent with the perspective that interspecific competition can be a strong force setting range limits in nature. Dispersal behavior remains poorly understood in nature, and our theoretical findings point to rewarding lines of enquiry. Experiments measuring movements of individuals near range borders, adaptive or maladaptive effects of gene flow to peripheral populations, and phenotype-dependent competitive interactions would be useful to test the predictions of our work. In addition, observational data could be used to verify the result we present from theory, that character displacement is more common when dispersal is random compared to when dispersal is adaptive. Future empirical work can thus test whether and how our findings about the role of dispersal behavior in shaping species’ range limits apply in nature.

Matching habitat choice is a phenotype-environment matching dispersal strategy (Box 1). In the family of models such as the one we used in our study, the environment is modeled through the optimum value that it imposes on individuals’ phenotype. This trait optimum is often assumed to be determined by abiotic factors. However, a habitat can be described by biotic factors as well. In particular, it can be argued that phenotype-dependent competition (as in our model) can be included into the description of the environment. In this case, competition will directly contribute to individuals’ dispersal decision and direction, rather than the indirect contribution through creating a diversifying selection and inducing character displacement as in our study. Noting the conflicting effects of phenotype-(abiotic) environment matching dispersal and phenotype-dependent competition (Box 1), the individuals’ dispersal behavior in such a generalized form of matching habitat choice would then require assessment of a trade-off between avoiding competition and matching the optimum phenotype. Such an assessment could depend on population density as well. Including all such factors in the dispersal strategy makes it a rather fitness-maximizing (ideal free) optimal strategy, understanding the effects of which on coevolution of range borders can be an interesting direction of future research. Based on our analyses, we hypothesize that the formation of range borders by interspecific competition will still be robust to such an optimal dispersal behavior. However, the interactions between random and optimal dispersal and character displacement will likely converge to a different equilibrium state. An extension of our model that incorporates such a generalized habitat choice mechanism can help test our hypothesis and identify the impacts of optimal dispersal on the sharpness of the borders and extent of character displacement.

Finally, it is worth noting that adaptation to environments can also occur through other processes such as phenotypic plasticity and environment adjustment (niche construction). Understanding how these processes evolve independently or together, or along with matching habitat choice, and how the operation of each or a combination of them affects the coevolution of species’ range borders, is an important and challenging topic of research. The preliminary model-based studies by Edelaar et al. (2017); Goldberg and Price (2022); Scheiner et al. (2022), for example, can help the future efforts.

## Funding acknowledgment

FS was supported in part by NSF grant PHY 2146260 to Daniel Weissman and in part by grants from the NSF (DMS-2235451) and Simons Foundation (MPS-NITMB-00005320) to the NSF-Simons National Institute for Theory and Mathematics in Biology (NITMB). BGF was supported in part by Packard Foundation award 2024-77385.

## S1. Model Description

Our theoretical study in the present work is based on a comprehensive mathematical model which we build as an immediate extension to a sequence of previously developed evolutionary models. The extension we perform does not require additional mathematical derivations and is accomplished by combining the results developed by Shirani and Miller (2022, 2025) and Shirani and Freeman (2025). Therefore, below we first briefly state some historical remarks on the development of the model and then provide its main equations, which are the equations we numerically solve to obtain the computational results of our work. To better understand the major eco-evolutionary processes that are incorporated into the model we then describe each of the underlying components of the model. We also specify the key assumptions that are used in building each component and deriving the final equations. We refer the reader to the work of Shirani and Miller (2022, 2025) and Shirani and Freeman (2025) for further details. We note that the notations we use to present the model are the same as those used by Shirani and Miller (2022, 2025). These notations differ from the notations used by the proceeding models, to some extent. These notational differences are listed in Table 2 in Section S1.6 below.

### S1.1 Historical background of the development of the model

The mathematical model we use in our study has evolved through a sequence of developments, starting form the model proposed by Pease et al. (1989). In a seminal work aimed to test the “genetic swamping” hypothesis (Haldane, 1956; Mayr, 1963) as a cause of species’ range limits, Kirkpatrick and Barton (1997) used the preliminary results of Pease et al. (1989) to develop a model of a species’ range evolution over an environmental gradient in a fitness-related trait optimum. Using a system of partial differential equations, Kirkpatrick and Barton’s model represents the joint evolution of the population density and the mean value of a quantitative phenotypic trait for a single (solitary) species in a one-dimensional geographic space. The species’ dispersal in Kirkpatrick and Barton’s model is assumed to be in the form of diffusive (random) migrations, and the population growth rate is assumed to be logistic. The species’ adaptation to new environments is through the force of natural selection, which penalizes phenotypes that differ from the environment’s optimum phenotype. Importantly, the trait variance in Kirkpatrick and Barton’s model is assumed to remain constant in space and time. This unrealistic assumption was then relaxed by Barton (2001), who extended the model to further allow for the evolution of the trait variance.

The next important contribution to the development of models was made by Case and Taper (2000). In their influential work, Case and Taper extended the single-species model of Kirkpatrick and Barton to a community of competitively interacting species, where the competition between the individuals depended on their phenotype. However, trait variance in Case and Taper’s model was still assumed to be constant in space and time; an assumption they called “tenuous”. This assumption was then relaxed by Shirani and Miller (2022), who extended Case and Taper’s model to higher dimensional geographic spaces and allowed for the evolution of trait variance jointly with the population density and trait mean of each species within the community. Shirani and Freeman (2025) subsequently added Allee effects to the model, which allowed for removing some biologically unrealistic solutions of the model that would arise in certain evolutionary regimes.

The individuals’ dispersal (migration) in all the models described above is assumed to be random. In a recent development aimed to study the effects of matching habitat choice on range evolution of a species, Shirani and Miller (2025) worked on a single-species version of their model and additionally incorporated phenotype-optimal dispersal, a special type of matching habitat choice in which individuals follow the direction of environmental gradient in the trait optimum to settle in the habitat best suited for their phenotype. In the present work, we further extend the single-species model developed by Shirani and Miller (2025) to a community of competitively interacting species with phenotype-optimal dispersal. We also incorporate Allee effects. Since each individual’s optimal dispersal in Shirani and Miller’s model depends only on its own phenotype and not on other individuals’ phenotypes, the extension appears to be immediate and does not require further mathematical derivations. In Section S1.2 below, we provide the final equations of the model obtained by combining the equations developed by Shirani and Miller (2022, 2025) and Shirani and Freeman (2025). Since the presentation of the model with the community structure naturally appears to be complicated, an intuitive interpretation of each and every term in the equations is hard to establish. For the single-species case, the detailed interpretations of each terms in the model are available in Section 3 of the work by Shirani and Miller (2025). Such interpretations can provide helpful mechanistic insight into the key eco-evolutionary processes that shape the range dynamics predicted by the model.

### S1.2 Model equations

We specify the model for a community of N species, although throughout the present work we only focus on the coevolution of borders between two species (N = 2). The species’ habitat is modeled by an open rectangle Ω ⊂ ℝ^m^ in an m-dimensional geographic space, m ∈ {1, 2, 3}. In the present work we always consider a one-dimensional habitat, m = 1. The evolution time horizon is denoted by *T*, where *T* can take any positive values and is set to be sufficiently large based on the specifics of each simulation. Each individual within each species is represented by a quantitative phenotypic trait such as body size. The environment imposes an optimum value for this quantitative trait, denoted by Q, which we typically assume changes linearly with *x* over the habitat Ω.

At every geographic location *x* = (*x*_1_, …, *x*_m_) ∈ Ω and time *t* ∈ [0, *T*], the population density of the *i*th species, *i* ∈ {1, … N}, is denoted by *n*_*i*_(*x, t*), the mean value of the trait within the *i*th species is denoted by *q*_*i*_(*x, t*), and the variance of the trait is denoted by *v*_*i*_(*x, t*). The coevolution of the species’ ranges in the model is represented by a system of partial differential equations that govern the joint evolution of these three population quantities for each species. In the equations of the model given below, for brevity we define a vector *u* := (*n*_1_, *q*_1_, *v*_1_, …, *n*_N_, *q*_N_, *v*_N_) that contains all these state variables.

The formulations of the eco-evolutionary processes incorporated into the model, as described in Section S1.5 below, also include several parameters whose definitions and biologically plausible ranges of values are given in Table 1. Further details of these parameters and their units are provided in Section S1.3. Note that in Table 1, the parameter D_*i*_(*x*) is an m × m matrix (D_*i*_(*x*) ∈ ℝ^m×m^), whereas the rest of the parameters are scalar-valued. Moreover, the parameters S, U, J_*i*_, V_*i*_, Π, and *κ* are assumed to be constant in space (independent of *x*), whereas D_*i*_, A_*i*_, K_*i*_, R_*i*_, and Q can be variable in space. All these model parameters may also vary in time, although their dependence on *t* is not explicitly shown in the equations (S1)–(S14) below.

To write the equations of the model, we denote the partial derivative with respect to *t* by ∂_*t*_, the gradient with respect to *x* by ∇_*x*_, the divergence with respect to *x* by div, the standard inner product in ℝ^m^ by 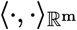, and the Euclidean norm in ℝ^m^ by 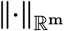.

**Table S1:**
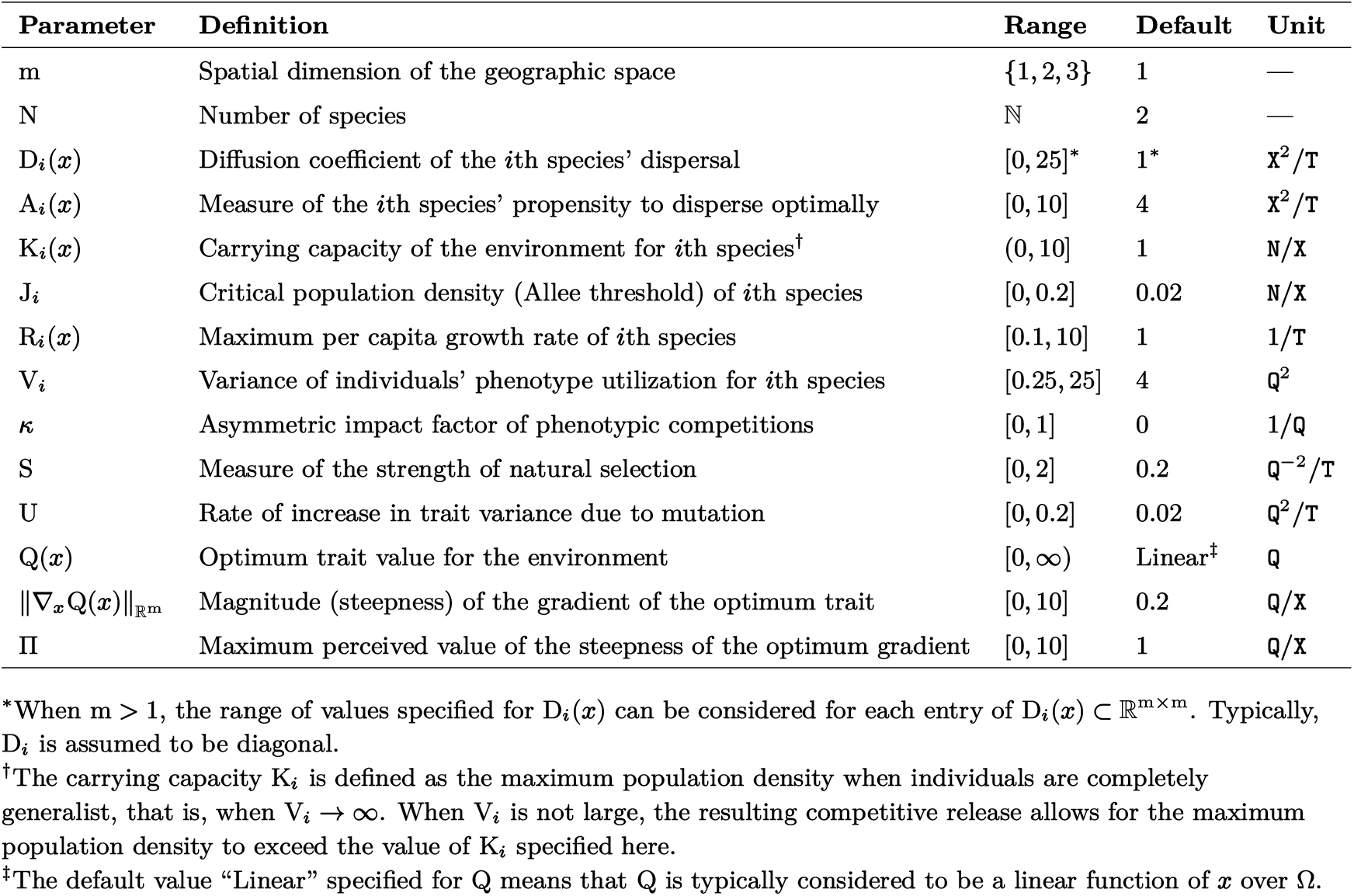
Definition and default values of the parameters of the model (S1)–(S14); (Shirani and Freeman, 2025; Shirani and Miller, 2022, 2025). The default values are used in all simulations presented in this work, unless otherwise is stated.

We start by writing the equation for the evolution of the population density *n*_*i*_ of the *i*th species, *i* = 1, …, N, as

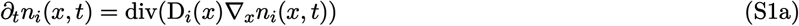

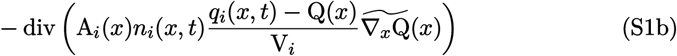

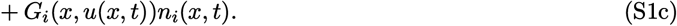

where *G*_*i*_(*x, u*) denotes the mean growth rate of the population given by equation (S4) below, and 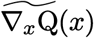 denotes individuals’ perceived environmental gradient (Section S1.5) given by equation (S14) below.

Likewise, the equation for the evolution of the trait mean *q*_*i*_ within the *i*th species is given as

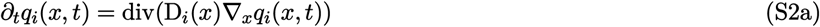

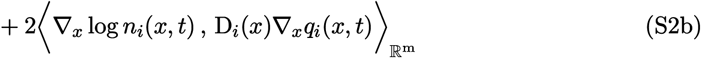

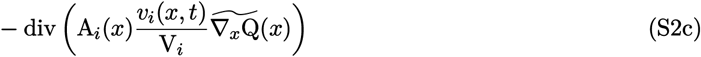

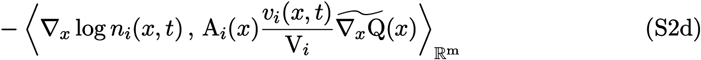

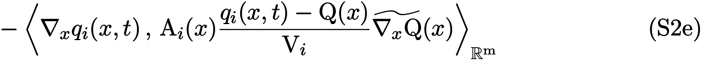

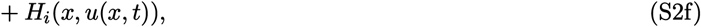

where *H*_*i*_(*x, u*) is defined by equation (S5) below.

Finally, the equation for the evolution of the trait variance *v*_*i*_ within the *i*th species is

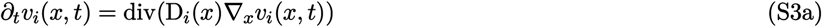

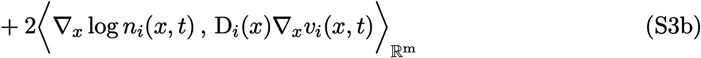

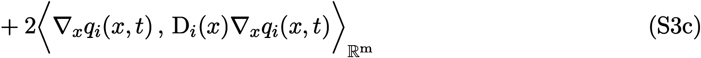

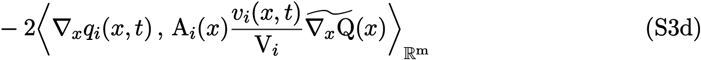

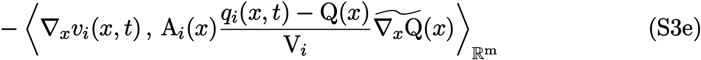

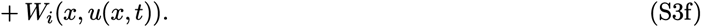

where *W*_*i*_(*x, u*) is defined by equation (S6) below.

The nonlinear mappings *G*_*i*_, *H*_*i*_, and *W*_*i*_ in (S1c), (S2f), and (S3f) are defined as

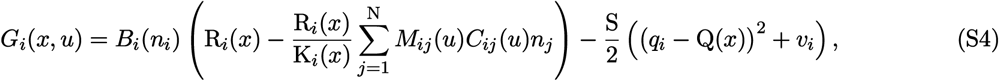

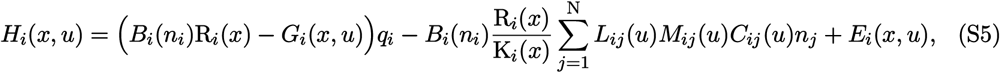

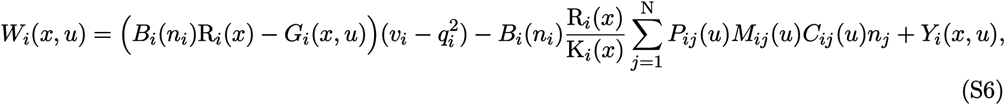

where, letting 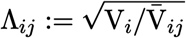 with 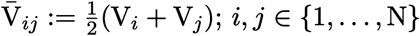,

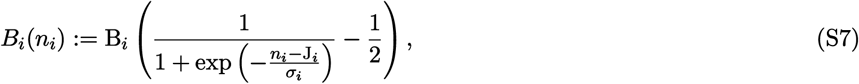

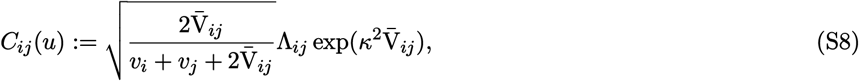

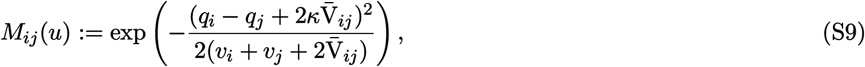

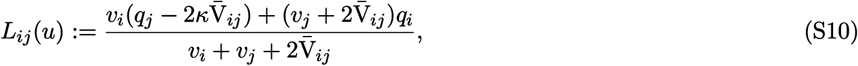

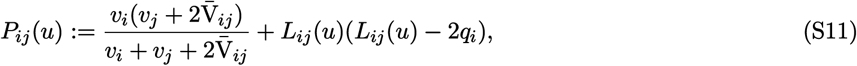

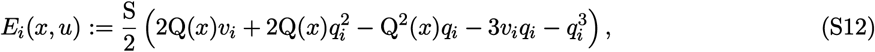

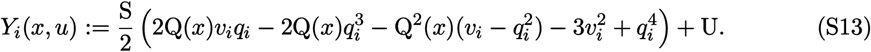

The nonlinear function *B*_*i*_(*n*_*i*_) given by (S7) adds Allee effects to the intrinsic population growth of the species, as described in Section S1.5 below. The key parameter in the formulation of *B*(*n*_*i*_) is the Allee threshold J_*i*_ as defined in Table 1. The parameter *σ*_*i*_ in (S7) determines how sharply the intrinsic growth rates increase to their maximum. The parameter B_*i*_ is adjusted such that the maximum value of *B*_*i*_(*n*_*i*_) becomes equal to 1, so that the maximum intrinsic growth rate of the species becomes equal to R_*i*_. For both species throughout our study we set J_*i*_ = 0.02 N*/*X and *σ*_*i*_ = 0.05 N*/*X. The parameter B_*i*_ is then calculated based on theses values of J_*i*_ and *σ*_*i*_. We obtain B_*i*_ = 2.6. See also Figure S1 in the work by Shirani and Freeman (2025).

Finally, the individuals’ perceived environmental gradient 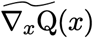 in (S2) and (S3) is defined as

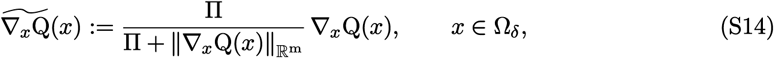

where Π is a constant that determines the maximum steepness of the environmental gradient that is perceived by the individuals, as defined in Table 1; see also Section S1.5 below. The smaller habitat Ω_*δ*_ specified in (S14) includes all points of Ω except those that are closer than a constant *δ* to the boundary of Ω. This is due to some technical reasons described by Shirani and Miller (2025). It is assumed that the individuals of the species are able to perceive the boundary of the habitat once they become sufficiently close to it (closer than the constant *δ*), and they avoid crossing the boundary. To compute the solutions of the model, the definition of 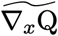 in (S14) must be extended to the entire habitat Ω. The formulation for such an extension in the general case of an m-dimensional habitat is given in Appendix A of the work by Shirani and Miller (2025). For the one-dimensional habitat Ω = (*a, b*) that we consider in the present work, the extension is performed as follows:

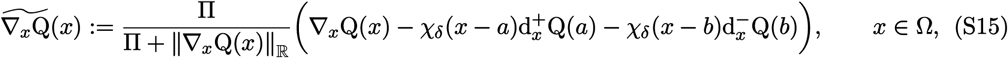

where 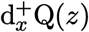 and 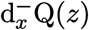 denote, respectively, the right-hand and left-hand derivatives of Q with respect to *x* evaluated at a point *z*, and

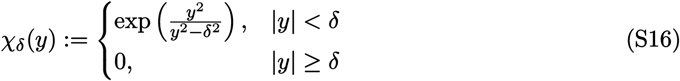

is a smooth cut-off function. We set the value of the parameter *δ* equal to 2 X.

Solving the equations of the model (S1)–(S14) also requires setting appropriate boundary conditions at habitat boundaries. In general, different boundary conditions such as reflecting, periodic, absorbing, or a combination of these conditions can be set based on the context of the problem under study and dimension of the geographic space. Some technical considerations for setting the boundary conditions are given in Remark 1 and Appendix A.5 of the work by Shirani and Miller (2022). For the one-dimensional habitat Ω = (*a, b*) that we consider in the present work, we assume no phenotype flux through the habitat boundary which, as discussed by Shirani and Miller (2025: Section 2.4), results in the homogeneous Neumann boundary conditions

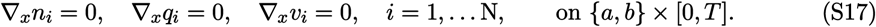

These conditions further imply that the habitat boundaries are reflecting. For example, if we conceptualize the habit to be representing the living environment of species of montane birds over an elevation-dependent temperature gradient, then the reflecting boundary condition at the high-elevation side of the habitat is equivalent to symmetrically (reflectively) expanding the habitat at the mountain top (Shirani and Freeman, 2025).

### S1.3 Model parameters: definitions, units, and ranges of values

Due to the complexity of the equations of the model (S1)–(S14), analytical studies of the behavior of the model are rather impractical. Numerical analyses can result in biologically meaningful predictions provided that they are performed with carefully chosen biologically plausible parameter values. This further requires deliberate choices of units for the demographic, ecological, and evolutionary quantities present in the model. An extensive discussion has been provided by Shirani and Miller (2022: Section 3) on reasonable choices of units that give sufficient generality for the model, as well as plausible ranges of values that the parameters can take based on such units. The discussion has been extended by Shirani and Miller (2025: Section 2.5) to further include the parameters of phenotype-optimal dispersal. Here, we only describe the choices of units as needed for understanding the values given in Table 1. We also provide some comments on the values of a key parameter of the model: the steepness of the gradient in environmental trait optimum.

To specify units for the physical quantities of the model, one of the species in the community is first chosen as a *representative species*, for example, the one which is best adapted to the environment or has the widest niche. The units are then specified based on measurements of basic quantities within the representative species. Specifically, the unit of time, denoted by T, is set to be equal to the mean generation time of the representative species. For the one-dimensional habitat that we consider in the present work, the unit of space, denoted by X, is chosen such that the diffusion coefficient D of the representative population becomes unity. That is, we set 1 × to be equal to the root mean square of the dispersal distance of the population in 1 T, divided by 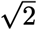. Estimates of the random component of the species’ dispersal can be obtained, for example, by measuring dispersal distance of a subpopulation of individuals that are well adapted to the environment at the core of the population and hence do not perceive a significant force to disperse optimally. Having set the unit of space, the unit of measurement for population abundances, N, is then chosen such that 1 N is equal to the carrying capacity of the environment for 1 × unit of habitat length (or, in general, 1 X^m^ unit of habitat volume). Note that this results in the default value of the carrying capacity for the representative population becoming unity. Finally, letting Q denote the unit of measurement for the quantitative trait, we set 1 Q to be equal to one standard deviation of the trait values at the core of the representative population. A toy example is provided by Shirani and Freeman (2025) to describe how empirical measurements based on other choices of units can be converted to estimates based on the units we described above.

The steepness of environmental gradients in trait optima is a key factor in the evolution of phenotype-dependent dispersal strategies such as matching habitat choice (Shirani and Miller, 2025), as well as the contribution of gene flow to the coevolution of species’ borders. Nevertheless, estimates of this key parameter are hard to obtain, and the available estimates are often based on different choices of units which make steepness comparisons almost impossible. In addition to measurements of the optimum trait value at different geographic locations, estimates of the environmental gradient based on the choices of units described above further requires measurements of standard deviation of the trait values, the mean dispersal distance, and the mean generation time of the representative population. The discussion given by Shirani and Miller (2022: Sect. 3.2) is based on limited data, and suggests a plausible range of values for the steepness of the gradient to be approximately between 0 Q*/*X and 2 Q*/*X. The values close to 10 Q*/*X, as listed in Table 1, can be used to model physical barriers. Finding more accurate estimates for a broad range of species— based on choices of units which provide sufficient generality for comparison purposes (such as the units described above)—will be an important contribution to future studies of species adaptation and range evolution in heterogeneous environments.

### S1.4 Model assumptions

The derivation of the equations of the model (S1)–(S14) relies on the following assumptions. The mathematical formulation associated with each assumption are given in Section S1.5 below.

i. Random dispersal of the individuals in each species is diffusive.
ii. An individual’s environmental *phenotypic potential energy* for phenotype-optimal dispersal is proportional to the square of the difference between its phenotype value and the environment’s optimum phenotype value.
iii. Nonlinear environmental selection for an optimal phenotype is stabilizing.
iv. The frequency of phenotype values within the species is normally distributed at every occupied point in space and for all time.
v. In the absence of selection, the intrinsic growth rate of the individuals follows a logistic growth with Allee effects and phenotype-dependent competition. The carrying capacity and maximum growth rate of the individuals are independent of their phenotype.
vi. Environmental resources vary continuously along a resource axis, and the resource utilization of each individual within each species follows a normal distribution. The resource axis can be identified by the phenotype axis through a one-to-one mapping. The individuals’ *phenotype utilization distribution*, obtained by mapping the resource utilization distributions, remains normal.
vii. The strength of intraspecific competition between individuals is determined by the overlap between their phenotype utilization curves, and can be asymmetric.
viii. The probability of mutational changes from one phenotype to another phenotype depends on the difference between the phenotypes, and is assumed to be normally distributed with zero mean and constant variance.
ix. For simplicity of model derivation, phenotypic variability is assumed to be predominantly genetic, and genetic variation is assumed to be predominantly additive. That is, H^2^ ≈ h^2^ ≈ 1, where H^2^ and h^2^ denote broad- and narrow-sense heritability, respectively.

### S1.5 Model components and the underlying eco-evolutionary processes

The equations of the model (S1)–(S14) are derived from a basic evolutionary equation that specifies, over a small interval of time, changes in population density of individuals with phenotype *p* ∈ ℝ within each species. To present this equation, let *φ*_*i*_(*x, t, p*) denote the relative frequency of phenotype value *p* ∈ ℝ in the *i*th population, at a location *x* ∈ Ω and time *t* ∈ [0, *T*]. Then *n*_*i*_(*x, t*)*φ*_*i*_(*x, t, p*) gives the population density of individuals with phenotype *p* in the *i*th population. The model assumes that changes in *n*_*i*_(*x, t*)*φ*_*i*_(*x, t, p*) over a small time interval of length *τ* → 0 results from the contribution of four major factors, as:

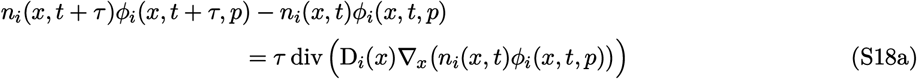

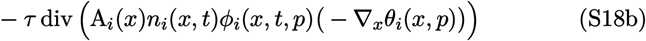

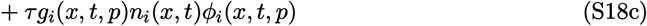

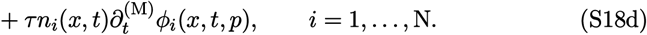

The term (S18a) represents diffusive (random) dispersal of individuals to and from neighboring locations. The term (S18b) represents directed (optimal) dispersal of individuals in the direction that gives them maximum environmental match. The individuals’ propensity A_*i*_ and perceived force −∇_*x*_*θ*_*i*_ in (S18b) are described below. The term (S18c) models the intrinsic population growth, where the intrinsic growth rate is denoted by *g*_*i*_(*x, t, p*) and is described below. Finally, the term (S18d) represents mutational changes in the relative frequency of *p* in the *i*th population, where the rate of mutational changes is denoted by 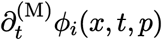 and is described below.

#### Intrinsic growth rates

A key factor that differentiates the density of individuals with different phenotypes is their intrinsic growth rate, modeled as

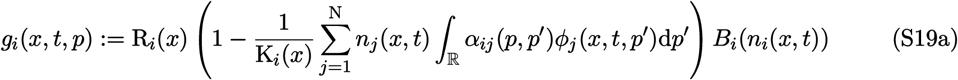

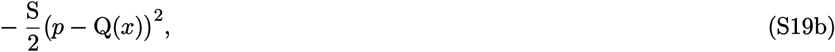

for the *i*th species in the community of N species. Parameter R_*i*_ denotes the maximum per capita growth rate of the species, K_*i*_ denotes the carrying capacity of the environment for the species when the species’ individuals are completely generalist (see below), S denotes the strength of stabilizing selection, and Q(*x*) denotes the environment’s optimal trait value at location *x* (see also Table 1). In the absence of the term *B*_*i*_(*n*_*i*_), (S19a) models a logistic growth, wherein the maximum growth rate R_*i*_ of the *i*th population occurs when all population densities *n*_*j*_, *j* = 1, …, N, go to zero. The carrying capacity and maximum growth rate of individuals are assumed to be independent of their phenotype (Assumption (v)). In the presence of *B*_*i*_(*n*_*i*_), defined by (S7), the population experiences Allee effects, so that it has negative growth rate when its density is lower than the Allee threshold J_*i*_. The convolution term in (S19a) models the effects of phenotypic competition between the individuals as described below. The term (S19b) incorporates the effects of directional and stabilizing natural selection on individuals with phenotype *p*. It penalizes phenotypes which are different from the environmental optimal value Q(*x*). This penalizing effect is stronger for larger values of S.

#### Competition kernels

The competition kernel *α*_*ij*_(*p, p*′) in (S19a) represents the strength of per capita effects of individuals with phenotype *p*′ in the *j*th species on the frequency of individuals with phenotype *p* in the *i*th species. Using the MacArthur-Levins overlap formula between *phenotype utilization curves* of each species (Shirani and Miller, 2022: Appendix A.2), the competition kernel is given as

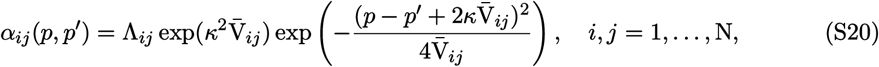

where 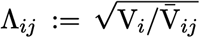 with 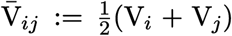. Parameter *κ* is used to model asymmetry in phenotypic competitions. Throughout this work, however, we always assume symmetric competition between phenotypes, with *κ* = 0. Parameter V_*i*_ denotes the variance of *phenotype utilization distributions*, which constitute the phenotype utilization curves and are assumed to be normal,

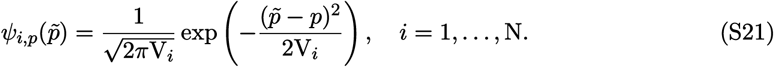

The phenotype utilization distribution *ψ*_*i,p*_ of individuals with phenotype *p* in the *i*th population are obtained from their presumed normally distributed resource utilization distribution, after identification of the resource axis with the phenotype axis (Shirani and Miller, 2022: Appendix A.2). The distribution function 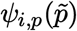 can be interpreted as a function that gives the probability density that an individual with phenotype *p* in *i*th species will utilize a unit of resource that is expected to be mostly utilized by (is most favorable for) an individual with phenotype 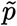. In trait-based niche quantification frameworks (Ackerly and Cornwell, 2007; Violle and Jiang, 2009) the variance of phenotype utilization distributions can be used to quantify within-phenotype component of a species’ niche breadth. Moreover, the resource-phenotype identification used to construct phenotype utilization distributions can be conceptualized to represent the empirical relationship between functional response traits and environments. This interpretation inspires the choice of the phenotypic potential energy function for optimal dispersal, as described below.

#### Carrying capacity

The phenotype-dependence of the competition in the model creates an ambiguity in the definition of K_*i*_, which needs to be clarified (Shirani and Freeman, 2025). As stated above, K_*i*_ denotes the maximum population density of the *i*th species when the species’ individuals are “completely generalist”, that is, when V_*i*_ → ∞, *i* = 1, … N. This is because *α*_*ij*_(*p, p*′) → 1 in (S20) when V_*ij*_ → ∞, regardless of the values of *p* and *p*′. Setting *α*_*ij*_(*p, p*′) = 1 in (S19) verifies that completely generalist species with density *n*_*i*_ *>* K_*i*_ will have negative intrinsic growth, hence the population density of such species will be bounded by K_*i*_. However, when individuals are specialized in utilizing resources, they become partially released from competition, meaning that *α*_*ij*_(*p, p*′) *<* 1. In this case, *g*_*i*_ can take positive values for some *n*_*i*_ *>* K_*i*_. That is, specialized species can grow to population densities greater than K_*i*_; see, for example, the curves of range expansion wave amplitudes given by Shirani and Miller (2025: Fig. 2b). Therefore, even though K_*i*_ is phenotype-independent, the carrying capacity of species—defined generally as species’ maximum population density—depends both on the distribution of phenotypes and individuals’ phenotype-based resource utilization.

#### Changes due to mutation

The rate of mutational changes in the frequency of phenotypes in the *i*th population is modeled as

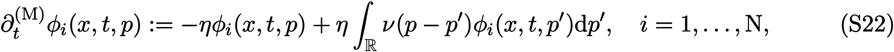

where *η* ≥ 0 is the mutation rate per capita per generation. Following Assumption (viii), the mutation kernel *ν*(*δp*) denotes the probability density that by mutation a phenotype *p* changes to a phenotype *p*′ = *p* + *δp*. The first term in (S22) incorporates the reduction rate in the frequency of phenotype *p* within the *i*th species due to mutation to other phenotype values. The second term incorporates the growth rate in the frequency of phenotype *p*, due to mutations from other phenotype values to *p*.

#### Phenotype-optimal dispersal

The phenotype-optimal dispersal in the *i*th species is modeled by the advection term (S18b), which is analogous to the drift component of drift-diffusion models of particles flowing in a fluid. The *propensity* of the *i*th species’ individuals to disperse optimally, denoted by A_*i*_, is analogous to the *mobility* of the particles. The *phenotypic dispersal force* −∇_*x*_*θ*_*i*_(*x, p*) induced by individuals’ perceived *phenotypic potential energy θ*_*i*_(*x, p*) is analogous to the force acting on particles derived from an external potential energy. The following model is used for the phenotypic potential energy function of an individual with phenotype *p* in the *i*th species at a habitat location *x*,

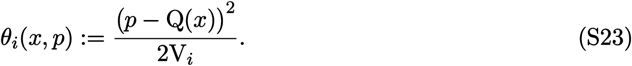

This potential energy function is proportional to a first-order approximation of the phenotype utilization distribution *ψ*_*i,p*_(Q(*x*)). It can therefore be interpreted as how well or poorly an individual with phenotype *p* can utilize resources at its current location *x*, which are most favorable to individuals with optimum phenotype Q(*x*). If the individual’s phenotype matches the optimum phenotype perfectly, then there is no phenotypically and environmentally induced force on the individual to disperse. If the individual’s phenotype differs significantly from the environment’s optimum—measured relative to the species’ phenotype utilization variance V_*i*_—then the individual perceives a high potential energy to disperse to habitat locations of better quality that match its phenotype. However, unless a sufficiently large gradient in the phenotypic potential energy is perceived by the individual, a high potential energy *θ*_*i*_(*x, p*) does not generate a significant dispersal force −∇_*x*_*θ*_*i*_(*x, p*) on the individual. Such a gradient is present if the environmental gradient ∇_*x*_Q is sufficiently steep and the individual is sufficiently sensitive to it.

The phenotypic potential energy (S23) gives an optimal dispersal force term −∇_*x*_*θ*_*i*_ as

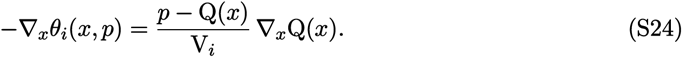

If the magnitude of the environmental gradient 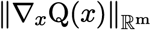 is zero (where 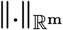 denotes the Euclidean norm in ℝ^m^) or is perceived as equal to zero due to insensitivity of an individual to the gradient, then the perceived directed dispersal force on the individual is zero—regardless of the presence of a phenotype-environment mismatch *p*≠ Q(*x*). In this case the individual only disperses randomly, due to the diffusion term (S18a). When 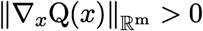, an individual whose phenotype *p* does not perfectly match the optimum phenotype will perceive a force pushing it to disperse optimally. If *p >* Q(*x*) the optimal dispersal will be in the direction of the environmental gradient, and if *p <* Q(*x*), the optimal match will be in the direction opposite to the environmental gradient.

A biological organism may not develop a perception of the phenotypic dispersal force (S24) in its exact mathematical sense. However, to make the modeling mathematically feasible, the phenotypic potential (*p* − Q)*/*V_*i*_ part of the dispersal force is assumed to be perceived by the individuals exactly. However, a biologically more reasonable perception of the environmental gradient, denoted by 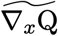 and formulated by (S14), is used for the gradient part. That is, the individual’s *perceived dispersal force* for phenotype-optimal dispersal,

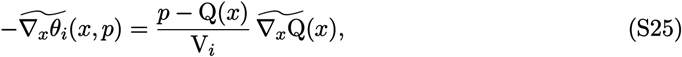

is substituted into (S18b) for the dispersal force term. The constant Π in the formulation of the perceived gradient (S14) determines the maximum perceived magnitude of the gradient. When 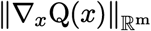 is much smaller than Π, the perceived gradient approximately equals the actual gradient. When 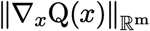 is much larger than Π, the magnitude of the perceived gradient approximately saturates to the maximum value Π. The direction of the perceived gradient, however, will always be the same as the direction of the actual gradient. The perceived gradient 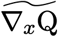 can be further interpreted, in some sense, as the sensitivity of the individuals’ dispersal force to changes in habitat quality. See Remark 3 of the work by Shirani and Miller (2025) for further details.

**Table S2:** Notational comparisons between the model presented in this work and Case & Taper’s model (2000). The presence of the enumeration index *i* in a variable implies that the variable can take a different value for each species.

| Parameter/variable/function description | Present work | Case & Taper’s (2000) |
| --- | --- | --- |
| The random variable denoting phenotype values | $p$ | $z$ |
| Population density | $n_i$ | $N_i$ |
| Trait mean | $q_i$ | $\bar{z}_i$ |
| Trait variance | $v_i$ | $V_p$ |
| Per capita intrinsic growth rate | $g_i$ | $w_i$ |
| Mean intrinsic growth rate | $G_i$ | $\bar{w}_i$ |
| Competition kernel | $\alpha_{ij}$ | $\alpha$ |
| Relative frequency (distribution) of phenotypes | $\phi_i$ | $p_i$ |
| Mutational changes in frequency of phenotypes | $\partial_t^{(M)}\phi_i$ | — |
| Diffusion coefficient of species’ dispersal | $D_i$ | $D$ |
| Measure of Species’ propensity to disperse optimally | $A_i$ | — |
| Carrying capacity of the environment | $K_i$ | $K$ |
| Critical population density (Allee threshold) | $J_i$ | — |
| Maximum per capita growth rate | $R_i$ | $r_i$ |
| Variance of individuals’ phenotype utilization | $V_i$ | $V_u$ |
| Asymmetric impact factor of phenotypic competitions | $\kappa$ | $\kappa$ |
| Measure of the strength of natural selection | $S$ | $1/V_s$ |
| Rate of increase in trait variance due to mutation | $U$ | — |
| Optimal trait value for the environment | $Q$ | $\theta$ |

### S1.6 Notational differences with preceding models

Compared with the notations used in the models of Kirkpatrick & Barton (1997) and Case & Taper (2000), some notational changes have been made here in presenting the equations and the underlying components of the model. These notational changes are made for several reasons, namely, to distinguish model variables from model parameters and mappings of model variables, to avoid conflicts with commonly used notations in mathematical analysis of PDEs, and to allow for further notational considerations in future extensions and analyses of the model. To facilitate comparisons with the preceding models, in Table 2 we provide the list of model parameters and variables with the notations used in the present work (as well as in Shirani and Freeman (2025); Shirani and Miller (2022, 2025)) and those used in Case and Taper (2000) (as well as in Kirkpatrick and Barton (1997) and other works in this family of models).

## S2. Numerical Computations

### S2.1 Numerical methods

The numerical method used in the work by Shirani and Miller (2022, 2025) is further extended in the present work to obtain the simulation results for the range evolution of two competing species with phenotype-optimal dispersal, and for quantifying the effects of directed gene flow on coevolution of the species’ boundary. The method is based on an Alternating Direction Implicit (ADI) scheme with two stabilizing correction stages (Hundsdorfer, 2002). In each iteration of the scheme, instead of solving the nonlinear algebraic equations involved in the computations, the linearized version of these equations are solved. The linearization is performed by computing the associated Jacobian matrix of the system symbolically (by hand) so that the Jacobian matrix is exact. The iteration time steps are then made smaller to compensate for the linearization error. The first and second space derivatives are approximated using fourth-order centered differences. See Appendix B of the work by Shirani and Miller (2022) for further details.

In all simulations, we discretized the one-dimensional space (habitat) with a uniform mesh of size Δ*x* = 0.1 X. For the majority of the simulations in which we aimed to compute the equilibrium curves accurately, we used small time steps of length Δ*t* = 0.002 T. For simulations that did not require computation of fully accurate solutions, we chose longer time steps of Δ*t* = 0.005 T or Δ*t* = 0.01 T.

The numerical computations of the solutions of the model are challenging, mainly due to the terms ∇_*x*_ log *n*_*i*_ in the equations (S2) and (S3) of the model. These terms are undefined when the population densities *n*_*i*_ are zero. See further details in Section 4.1 of the work by Shirani and Miller (2022). Even if we initialize the populations such that at least an infinitesimal population density is initially present everywhere, there is still a chance of reaching numerical singularities if the density of a species undergoes a long density decline (or extinction) regime—especially in the presence of Allee effect. Similar to the approach taken by Shirani and Freeman (2025), we avoid this singularity problem by replacing the terms ∇_*x*_ log *n*_*i*_ with ∇_*x*_ log(*n*_*i*_ + *ϵ*), *ϵ >* 0. We set *ϵ* = 10^−5^, which works well in our simulations and does not result in any noticeable impact on the solutions compared with the solutions of the original equations.

### S2.2 Simulation initialization

To obtain the numerical results presented here and in the main text, we consider a one-dimensional habitat Ω = (0 X, 50 X) ⊂ ℝ with the reflecting boundary conditions (S17). We initially introduce the two populations allopatrically, the 1st population centered at *c*_1_ = 15 × and the 2nd population centered at *c*_2_ = 25 X. We set the initial population densities at *t* = 0 T as *n*_*i*_(*x*, 0) = 0.5*ϱ*(|*x* −*c*_*i*_ |), *i* = 1, 2, where *ϱ* is a bump function. We assume that the initial populations are perfectly adapted to the environment at their center, *q*_*i*_(*c*_*i*_, 0) = Q(*c*_*i*_), but poorly adapted at other habitat locations, ∇_*x*_*q*_*i*_(*x*, 0) = 0.5 ∇_*x*_Q(*x*) for all *x* ∈ Ω. Finally, we set the initial populations’ trait variance equal to *v*_*i*_(*x*, 0) = 1 Q^2^. The values of the model parameters used in each simulation are specified independently for each figure, based on the values given in Table 1.

### S2.3 Computing the contribution of dispersal components, selection, and competition to rate of changes in trait mean and population density

The contribution of each component of dispersal to the rates of changes in species’ trait mean, ∂_*t*_*q*_*i*_, and species’ population density, ∂_*t*_*n*_*i*_, *i* = 1, 2 are shown in Figures 2 of the main text (for 1st species) and Figure S1 here (for 2nd species). The contribution of the local effects of selection and competition to ∂_*t*_*q*_*i*_ and ∂_*t*_*n*_*i*_ are also shown in Figure 3 of the main text and Figure S3 for 1st species. For computing the dispersal contributions we use the terms (S1a), (S1b), and (S2a)–(S2e) in the model, which directly originate from the random and optimal dispersal of phenotypes as in (S18a) and (S18b). For computing the local contributions due to selection and competition we use the terms (S1c) and (S2f) in the model, which originate from the intrinsic growth rate of the population of phenotypes as in (S18c). These terms, along with (S3f), control the local dynamics of the populations.

Specifically, the curves in Figure 2a are computed as the sum of the terms (S2a) and (S2b) in the equations of the model, with *i* = 1. The curves in Figure 2b are computed as the sum of the terms (S2c)–(S2e), with *i* = 1. The contribution of the total dispersal to ∂_*t*_*q*_1_, shown in Figure 2c is computed as the sum of the curves in Figures 2a and 2b, that is, the sum of the terms (S2a)–(S2e), with *i* = 1. The curves in Figures S1a–S1c are computed similarly, using the same terms in model equations, but with *i* = 2. The curves of local effects in Figures 3b and S3b are computed using (S2f) with *i* = 1. The final curves of ∂_*t*_*q*_1_ in Figures 3c are computed as the sum of the curves in Figures 3a and 3b. Similarly, the final curves of ∂_*t*_*q*_1_ in Figure S3c are computed as the sum of the curves in Figures S2c and S3b.

For contribution of the dispersal terms to ∂_*t*_*n*_1_, the curves in Figure 2d are computed using (S1a) with *i* = 1, the curves in Figure 2e are computed using (S1b), and the curves in Figure 2f are computed as the sum of (S1a) and (S1b). Similarly, the curves of contributions to ∂_*t*_*n*_2_, shown in Figures S1d–S1f are computed using the same terms in model equations, but with *i* = 2. The curves of local effects in Figures 3e and S3e are computed using (S1c) with *i* = 1. The final curves of ∂_*t*_*n*_1_ in Figures 3f are computed as the sum of the curves in Figures 3d and 3e. Similarly, the final curves of ∂_*t*_*n*_1_ in Figure S3f are computed as the sum of the curves in Figures S2f and S3e.

## S3. Range Evolution of a Solitary Species

The range evolution of a solitary species using the reduction of our model to a single-species is extensively analyzed by Shirani and Miller (2022, 2025), as well as many of the preceding important works (Barton, 2001; Kirkpatrick and Barton, 1997; Polechová and Barton, 2015). Below we provide a short summary of the key results.

### S3.1 Range evolution under random dispersal

With random-only dispersal, Kirkpatrick and Barton (1997) show that genetic swamping can cause formation of a limited range when the environment is sufficiently steep. Moreover, they show that the population becomes extinct if the steepness of the gradient exceeds a critical value that depends on the strength of selection, dispersal coefficient, and maximum growth rate. However, when trait variance is allowed to evolve, Barton (2001) and Shirani and Miller (2022) show that the limited range regime predicted by Kirkpatrick and Barton (1997) in a linear environment does not exist. The population will either expand its range indefinitely or go extinct, depending on the steepness of the gradient. This is because, the trait variance is inflated by random gene flow as the steepness of the environment increases. As a results, the standing phenotypic load imposed by natural selection on the population growth rate increases with the steepness of the gradient. This leads to population extinction when the steepness of the gradient exceeds a critical value; see (Shirani and Miller, 2022: Sec. 4.2 and Fig. 4) and (Shirani and Miller, 2025: Fig. 2a). In the absence of Allee effect, the critical gradient steepness is given as

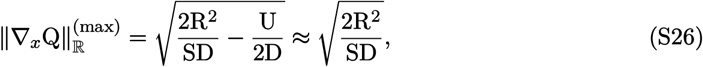

where the enumeration index *i* is dropped from the parameters, as we discuss only a single species here. The approximation in (S26) is made based on the fact that, typically, U ≪ 4R^2^*/*S. Note that (S26) implies that the critical gradient steepness decreases when R decreases or when S increases. This means that, slowly-growing species and species under strong selection are at higher risk of extinction at steep gradients. See (Shirani and Miller, 2022: Remark 4) and (Shirani and Miller, 2025: Remark 6) for more details.

The existence of the critical (extinction) gradient steepness described above also implies that steepening nonlinear gradients can limit species’ range (Bridle et al., 2019; Polechová, 2018; Polechová and Barton, 2015). This follows from the fact that gradient is a mathematical notion that is understood (defined) locally. A steepening nonlinear environmental gradient can be approximated by a piecewise linear gradient, such that the steepness of linear pieces increases along the environment. It then follows immediately from the predictions of the models (in linear environments) that the local population density over the linear pieces decreases as the steepness of the pieces increases. If the steepness of the steepening gradient eventually exceeds the critical extinction threshold predicted by the models, the local populations over exceedingly steep pieces will fail to service. This results in the formation of a range limit by the steepening gradient. Figures 10a and 10b of the work by Shirani and Miller (2022), for example, show how species’ range expansion stops at a steepening gradient.

### S3.2 Range evolution in the presence of matching habitat choice

Similar to the random-only dispersal case, our model predicts that in the presence of phenotype-optimal dispersal, the species’ range evolution in linear environments can have two different regimes: indefinite range expansion and complete extinction (Shirani and Miller, 2025). However, phenotype-optimal dispersal substantially increases the critical gradient steepness beyond which the species becomes extinct. This is because matching habitat choice substantially reduces local trait variance, and hence the phenotypic load imposed by selection on population growth (Shirani and Miller, 2025: Fig. 2a). Therefore, the species’ chance of survival at steep gradients is significantly enhanced by dispersing through matching habitat choice. Moreover, steepening nonlinear gradients should reach a much higher level of steepness in order to limit the range expansion of a species that disperses through matching habitat choice. For the majority of species in nature, such a steep gradient may not be realistically present or may just correspond to obvious physical barriers or habitat discontinuities.

In the absence of Allee effects, the critical gradient steepness (extinction steepness) can be computed by solving

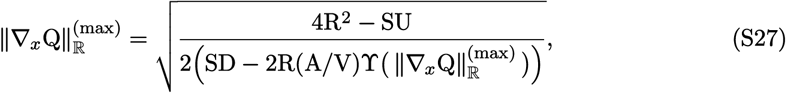

where 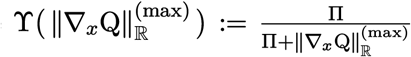. Similar to the random-only dispersal case, this critical steepness decreases as R decreases or as S increases. However, dispersing through matching habitat choice can still make the critical gradient steepness significantly greater than the gradients that realistically present in nature, even for (vulnerable) slowly-growing species under strong selection; see (Shirani and Miller, 2025: Fig. 3). This implies that, this class of vulnerable species may particularly benefit from evolving matching habitat choice, especially when the environment is fluctuating rapidly and frequently (Shirani and Miller, 2025: Figs. 6 and 7).

## S4. Supplementary Figures

**Figure S1:**
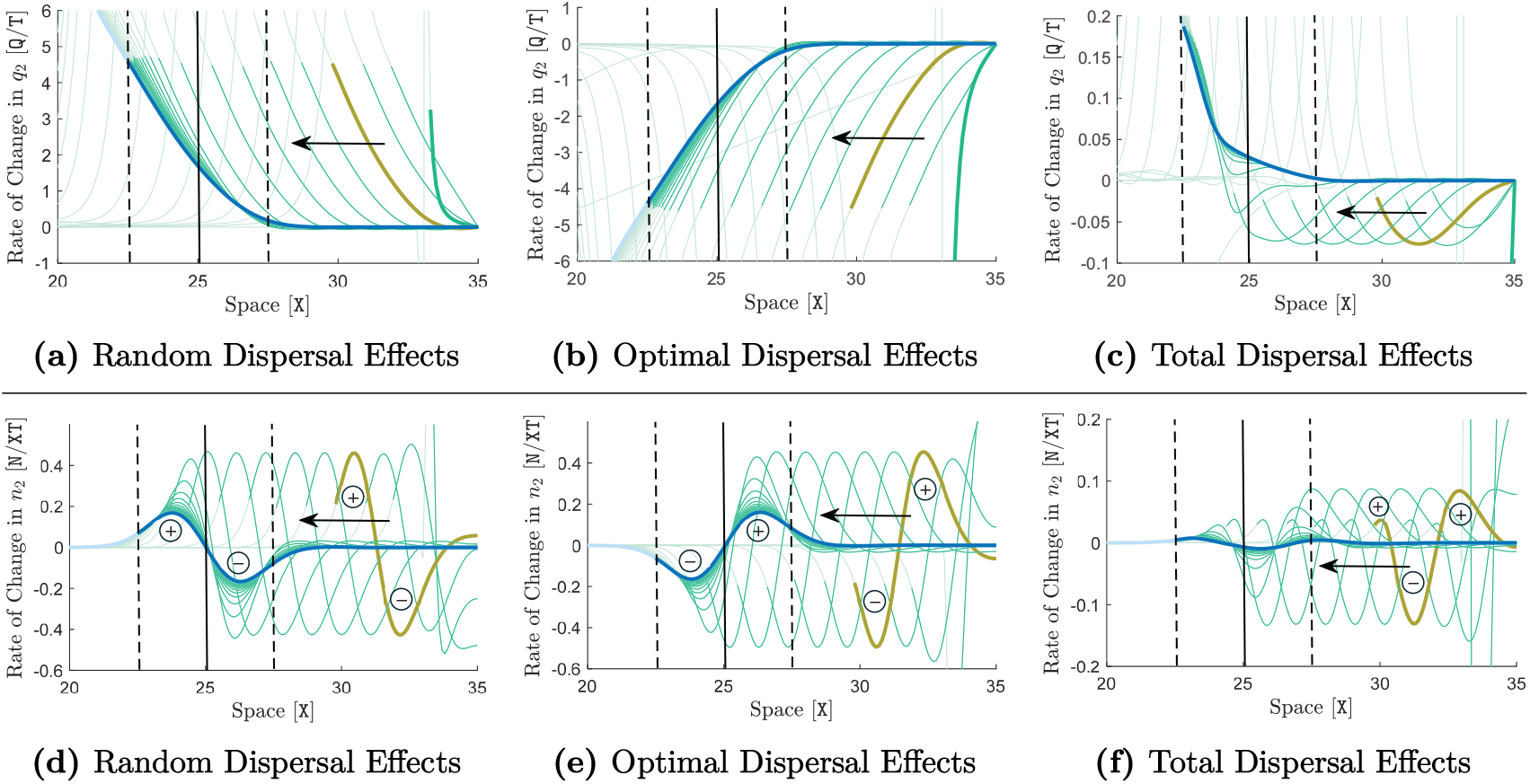
Effects of random and optimal dispersal on rates of changes in trait mean and population density. The results shown here are associated with the same analysis made to obtain the results shown in Figure 2 of the main text, but here we show the curves for the 2nd species. Specifically, the graphs are associated with the same simulation performed to obtain the results shown in the right panel of Figure 1, that is, with A_1_ = A_2_ = 10 X^2^*/*T and ∇_*x*_Q = 1.5 Q*/*X. The contributions of random dispersal, phenotype-optimal dispersal, and total dispersal to the rate of change of trait mean within the 2nd species, ∂_*t*_*q*_2_, are shown in graphs (a)–(c) in the upper panel. The contributions of these dispersal components to the rate of change of population density of the 2nd species, ∂_*t*_*n*_2_, are shown in the lower panel. The details of the computations associated with these contributions are described in supplementary Section S3.3. The curves in all graphs are shown only for the left half of the species’ range, where it meets and coevolves with the 1st species. In all graphs, curves are shown at every 4 T, and the parts of the curves which lie outside the species’ range (*n*_1_ *<* 0.02) are made transparent. The final equilibrium curves obtained (approximately) at the end of the simulation (*T* = 300 T) are highlighted in blue. The sample curves highlighted in olive are associated with the species’ range expansion regime before meeting the 1st species. The solid black lines indicate the center of the habitat where the interface between the two species (Figure 1b in the main text) is formed. The dashed lines indicate the boundaries of the region of sympatry formed at the interface.

**Figure S2:**
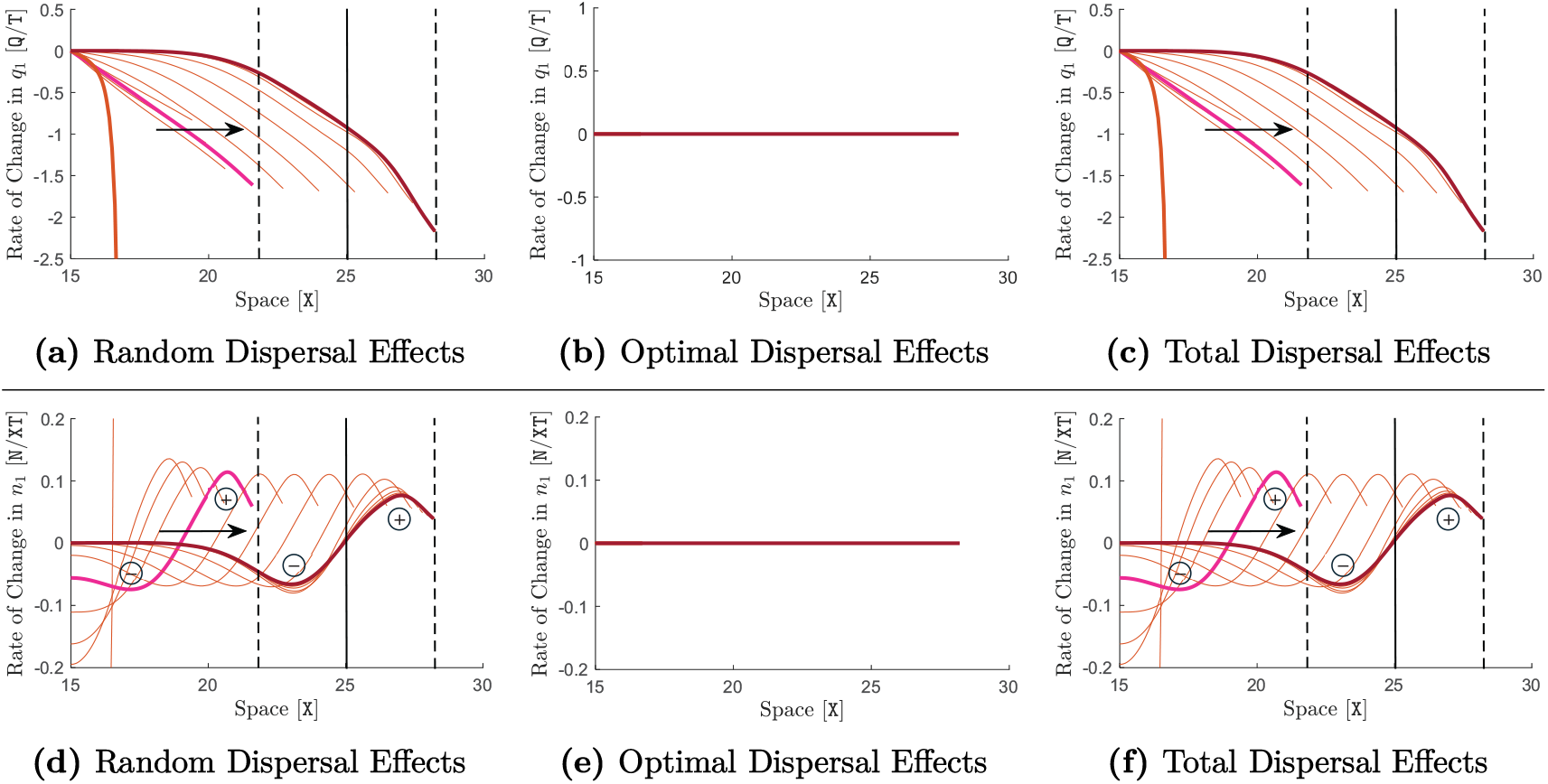
Effects of random-only dispersal on rates of change in trait mean and population density in the absence of phenotype-optimal dispersal. The results shown here are similar to those shown in Figure 2 of the main text, but here in the absence of phenotype-optimal dispersal (only with random dispersal). Specifically, the graphs are associated with the same simulation performed for the results shown in the left panel of Figure 1 in the main text, that is, with A_1_ = A_2_ = 0 X^2^*/*T and ∇_*x*_Q = 1.5 Q*/*X. Since phenotype-optimal dispersal is absent, its contribution to ∂_*t*_*q*_1_ and ∂_*t*_*n*_1_ is zero. The total contribution of dispersal to ∂_*t*_*q*_1_ and ∂_*t*_*n*_1_ is then equal to the contribution of random dispersal. The same description as in Figure 2 of the main text holds for the curve colors and the solid and dashed lines.

**Figure S3:**
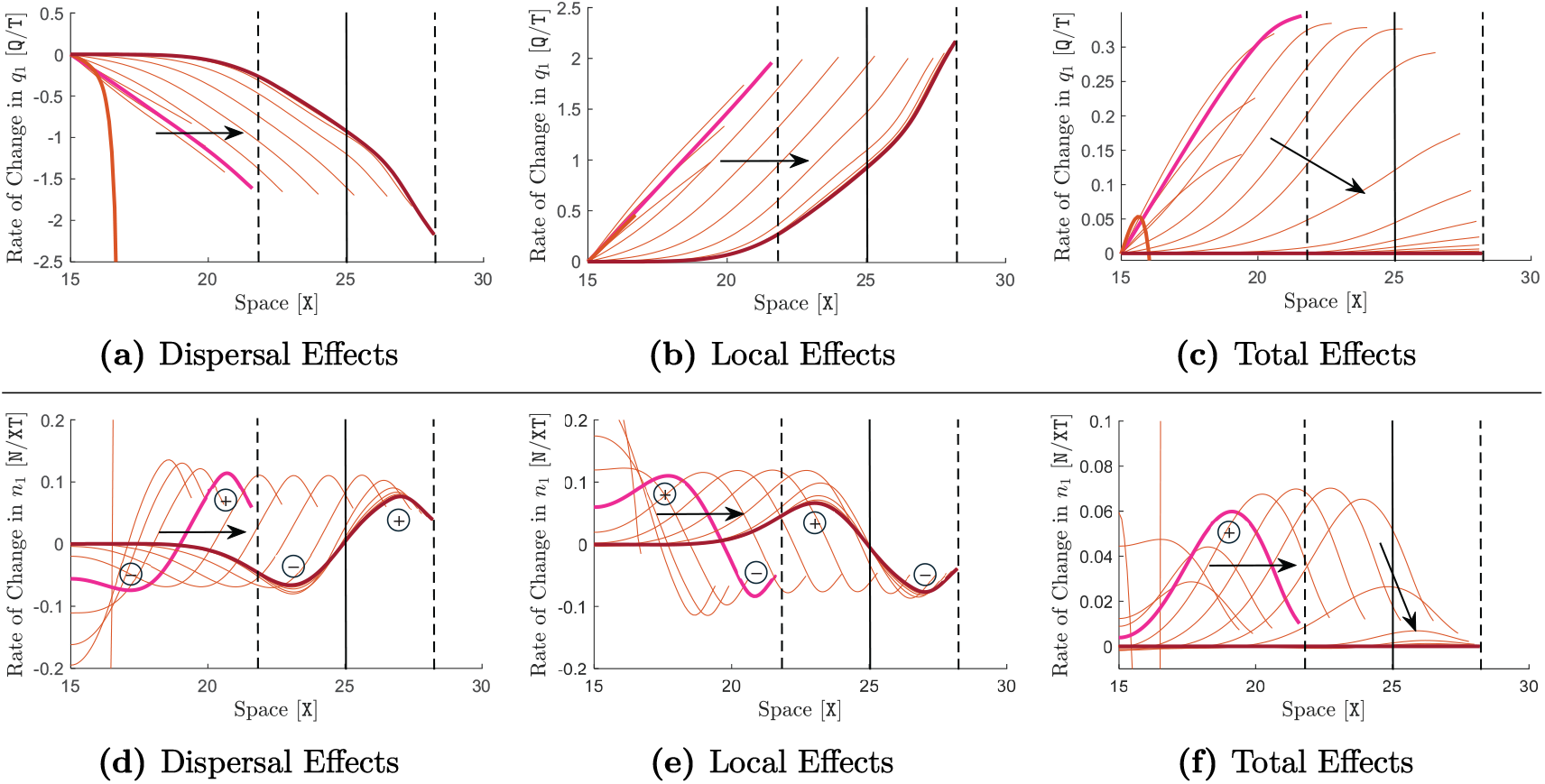
Effects of random-only dispersal, selection, and competition on rates of changes in trait mean and population density. The results shown here are similar to those shown in Figure 3 of the main text, but here in the absence of phenotype-optimal dispersal (only with random dispersal). Specifically, the graphs shown here complete the set of graphs shown in Figure S2 to demonstrate the effects of different components contributing to ∂_*t*_*q*_1_ (top panel) and ∂_*t*_*n*_1_ (bottom panel). As in Figure S2, the graphs are associated with the same simulation shown in the left panel of Figure 1 in the main text. Graphs (a) and (d) are the same graphs shown in Figures S2c and S2f, which are repeated here for simplicity of comparison. They show the contribution of random dispersal to ∂_*t*_*q*_1_ and ∂_*t*_*n*_1_, respectively. The local contributions of selection and competition to ∂_*t*_*q*_1_ and ∂_*t*_*n*_1_ are shown in (b) and (e), respectively. The details of the computations associated with these contributions are described in Section S2.3. Graphs (c) and (d) show the total effects, that is, they exactly show ∂_*t*_*q*_1_ and ∂_*t*_*n*_1_, respectively. The same description as in Figure 2 of the main text holds for the curve colors and the solid and dashed lines.

**Figure S4:**
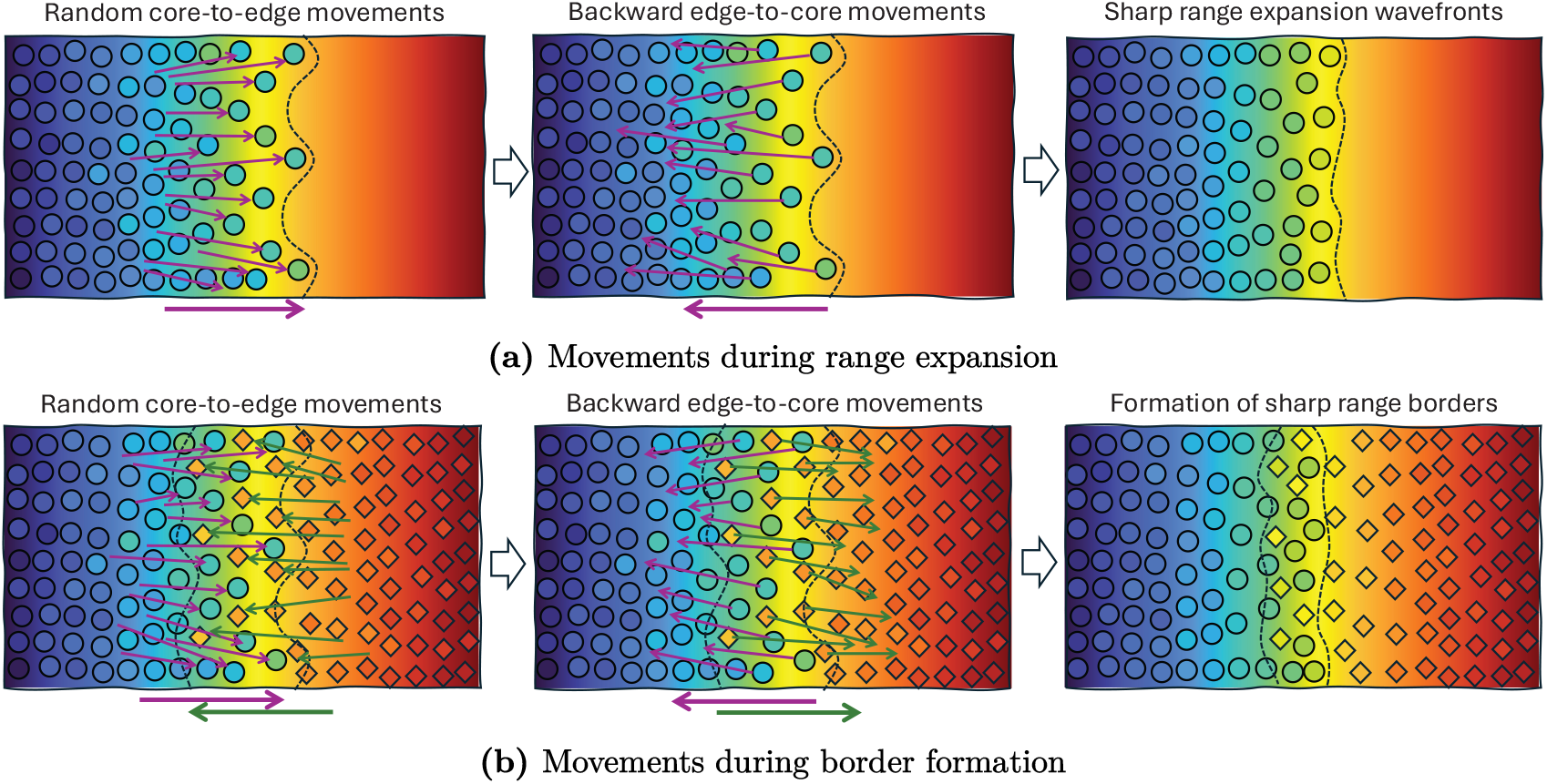
Schematic illustration of the movements caused by random and adaptive dispersal behaviors near the species’ range margins. In each graph, the predominantly core-to-edge movements caused by the random (diffusive) component of dispersal are shown on the left, and the the backward edge-to-core movements caused by matching habitat choice are shown in the middle. Although these different movement directions are shown separately here, for visualization clarity, they occur dynamically together during species’ range evolution. The background color gradients indicate the gradient in optimum phenotype. Individuals of the 1*st* species are shown by circles, and individuals of the 2nd species are shown by diamonds. The individuals’ color indicates their phenotype value. In (a), range expansion of a solitary species is shown. The individuals which disperse randomly from the range core to the range edge, possibly to explore new habitat, often face environments less suited for their phenotype. Following their matching habitat choice strategy, these individuals then move back to the core. These backward edge-to-core movements sharpen the range expansion wavefronts, illustrated in the diagrams on the right. Matching habitat choice results in strong adaptation (phenotype-environment match) except over a narrow region near the edge. Note that despite the sharpness of the wavefronts, the range edges shown by dashed lines keep moving forward and the species’ range expands continuously. In (b), the formation of range limits between two competitively interacting species is shown. Interspecific competition and maladaptive effects of random movements tend to induce a high degree of character displacement within the region of sympatry (between the dashed lines). This can be seen in the left diagram, through the distinct difference between the color of the individuals of the two species located between the dashed lines. However, matching habitat choice pushes the individuals whose phenotype differs significantly from the optimum to move back towards their population’s core. These backward movements (along with interspecific competition) eventually result in formation of sharp range limits (a narrow region of sympatry) and substantially reduced character displacement (relatively close colors).

